# The Muskox Way: Humans, plants and animals in northeast Greenland through the Holocene

**DOI:** 10.64898/2026.10.20.760410

**Authors:** Inger Greve Alsos, Marieke Beaulieu, Nicholas L. Balascio, Lucas Elliott, Raymond S. Bradley, Sergi Pla, Santiago Giralt, Luisa Deppe, Youri Lammers, Jamie Alumbaugh, Peter D. Heintzman, Love Dalén, Anders Götherström, Sandra Nogué, Bianca Perren, William J. D’Andrea, Frederik Fuuja Larsen, Christian Koch Madsen, Jostein Bakke

## Abstract

The Muskox Way hypothesis suggests that the first humans entering northeast Greenland from Arctic Canada followed and hunted muskox. Yet whether muskox arrived before people remains debated, as only two bones predate the Independence culture (4500–3900 cal BP). More broadly, the arrival of plants and animals, and how the three periods of human settlement related to ecosystem changes, remain poorly understood. We analysed sedimentary ancient DNA from lakes in Wandel Dal (Late Sommersø and Midsommersø) and Zackenberg, identifying 121 native plants and nine animal taxa. In Wandel Dal, the plant assemblage indicates a warmer climate than today about 8200-3900 years ago, followed by an abrupt change in flora and sedimentation rate at 3900 cal BP, especially at Midsommersø. Muskox DNA is present continuously at Midsommersø from 7400 cal BP, about three millennia before humans arrived. The Zackenberg record dates to 12,200 cal BP, with arctic forbs, grasses, ferns and crowberry in the earliest layers, willow from 8200 cal BP and birch from 7200 cal BP, but muskox only in the last 2500 years, the latter period coinciding with Greenlandic Dorset culture. The arrival of the Thule culture in 1400 CE coincides with a slight increase in plant richness at Wandel Dal and an increase in mammal detections. Reindeer occurred only in samples older than 6000 years, consistent with their absence from archaeological bone records. Overall, humans arrived long after the first megafauna and appear to have had limited impact on their populations, although cultural periods coincided with environmental changes.

**Significance:** Ancient DNA from lake sediments at three sites in northeast Greenland reveals the region’s most detailed ecological history, with 121 plant and nine animal taxa identified. Muskox reached the region about 7400 years ago, nearly 3000 years before human colonization, challenging the idea that people first entered Greenland by following their game. Muskox persisted throughout Independence culture occupation, suggesting that early hunting did not cause local extirpation. Around 3900 years ago, the end of this occupation coincided with environmental deterioration and major vegetation change. Greenlandic Dorset occupation coincided with vegetation change in Wandel Dal and the first detection of muskox at Zackenberg, showing that human settlement and ecosystem change were linked in time but not necessarily causally related.

## Introduction

Northeast Greenland (north of 74°N) represents the longest post-glacial colonization distance for Arctic biota and the northernmost prehistoric human settlements in the Arctic (1–3). The early “Muskox Way” hypothesis (4, 5) tried to explain the initial arrival of people by proposing that humans migrated into Greenland from Arctic Canada following muskox (*Ovibos moschatus*) as the climate warmed a hypothesis also supported by (6) (Fig. 1). In northeast Greenland, three distinct settlement phases were separated by millennia, raising further questions about how environmental change, particularly the availability of huntable animals, influenced the timing of human occupation. The use of the fossil record to address these questions is limited due to its fragmentary nature, owing largely to bone degradation and the rarity of fossil records outside of archeological settings (7). Advances in sedimentary ancient DNA (sedaDNA) analysis, which offer higher taxonomic resolution than pollen records and can detect mammalian presence even in the absence of bones (8–10), can shed new light on the post-glacial human-environmental history of this complex region.

**Fig. 1.**
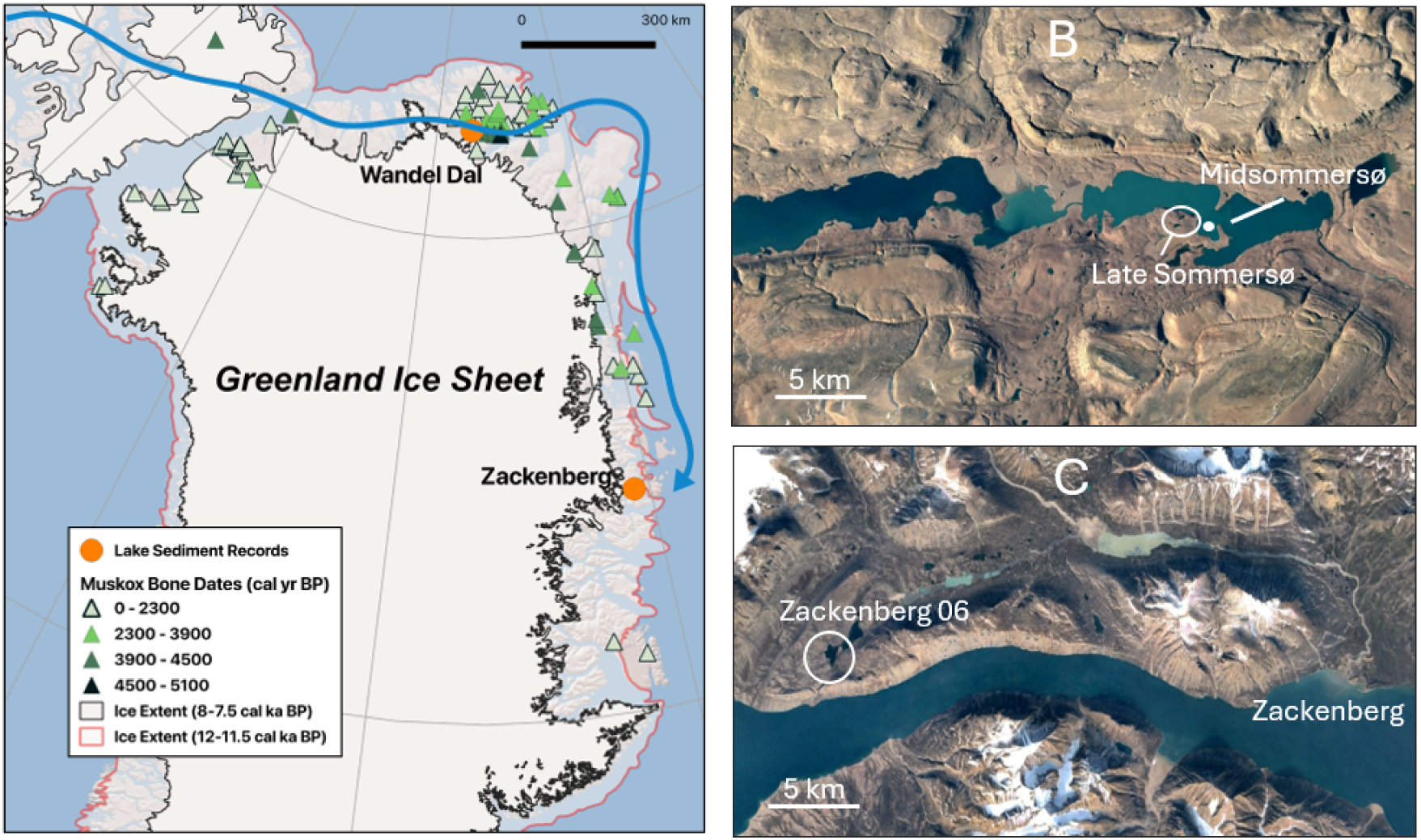
Study sites in Wandel Dal and Zackenberg in northeast Greenland. (A) The extent of the Greenlandic Ice Sheet over Greenland and the Innuitian Ice Sheet over Ellesmere Island at c. 12,000 cal BP (red line) and 8,000 cal BP (black line) according to PaleoGrIS version 1.0 using data from (22) and (23), respectively. Dated muskox bones are indicated, and the route that humans followed according to the Muskox Way hypothesis (4) is shown in blue. (B) The lakes “Late Sommersø” and Midsommersø in Wandel Dal. (B) The “Zackenberg 06” lake.

Although some ice-free areas in northeast Greenland persisted during the Last Glacial Maximum (LGM; ∼20,000 years ago), the majority of plants and all mammals colonized the region only after deglaciation (7, 11). The oldest post-glacial plant records consist of High- Arctic forbs and bryophytes and date to approximately ∼11,800 cal BP (calibrated calendar years Before Present), whereas shrubs and dwarf shrubs arrived 10800-6700 cal BP (11–13). Vertebrates arrived considerably later than plants, with the earliest scattered bird records dated to ∼7500 cal BP, Arctic char dated to the mid Holocene, and Arctic hare and Arctic lemming around 8000 and 6400 cal BP, the latter only based on dated droppings, whereas the vast majority of animal records are from archeological finds (7).

Archaeological assemblages in the region are from three cultural groups – the Independence (c. 4500-3900 cal BP), Greenlandic Dorset (c. 2800-2400 cal BP), and Thule (c. 1400 CE) – and consist of around ten thousand bone fragments (1, 3, 14). Muskox bones suggest dispersal from Ellesmere Island into Greenland via the Robeson Channel (15, 16). Although Campos et al. (15) argue for an almost synchronous entry of humans and muskox into northern Greenland around 4500 cal BP, but Jensen and Godfredsen (3) attribute this pattern to the predominance of Independence bone dates; indeed, two stray bones dated to ∼5100 cal BP (17) and ∼5400 cal BP (15) imply that muskox arrived at least a few hundred years prior to humans. A few dated bones suggest that muskox expanded southwards during or after the Greenlandic Dorset period, whereas more numerous dates from 1200 BP to the present indicate expansion to modern distribution (18).Antlers dates indicate that reindeer entered northern Greenland by ∼8800 cal BP, but their duration is uncertain because their scarcity in archaeological assemblages could reflect either local absence or hunting practices (6, 19, 20).

Wandel Dal in Inutoqqat Nunaat (Peary Land) has the highest density of archaeological records, whereas Zackenberg represents the southernmost extension of the Independence culture in northeast Greenland (1, 14, 16). Here we analyse sedaDNA from lake sediments in Midsommersø and “Late Sommersø” in Wandel Dal and one from “Zackenberg 06” (Fig. 1). We apply animal and plant metabarcoding to allow high-throughput detectability and taxonomic resolution, combined with shotgun metagenomics analyses of selected samples to authenticate the detections (21). Our aim is to 1) document the arrival of plants and animals to the region, and 2) explore changes in vegetation and how they relate to climate and herbivory. Based on our results, we discuss human occupation in relation to natural resources and environmental changes.

## Results

### Lithology and chronology

The Zackenberg, Midsommersø, and Late Sommersø cores span the past 12,200, 8200, and 7500 years, respectively. All three records show high Ca values in the Early Holocene, and Zackenberg and Midsommersø also show elevated sedimentation rates, together indicating warmer conditions, enhanced catchment runoff, and higher in-lake productivity before human arrival. A similar increase in sedimentation rates at Zackenberg and Late Sommersø during the Thule period may likewise reflect warmer conditions (Fig. 2, Supplementary Information 1, Supplementary Tables S1-S5).

**Fig. 2.**
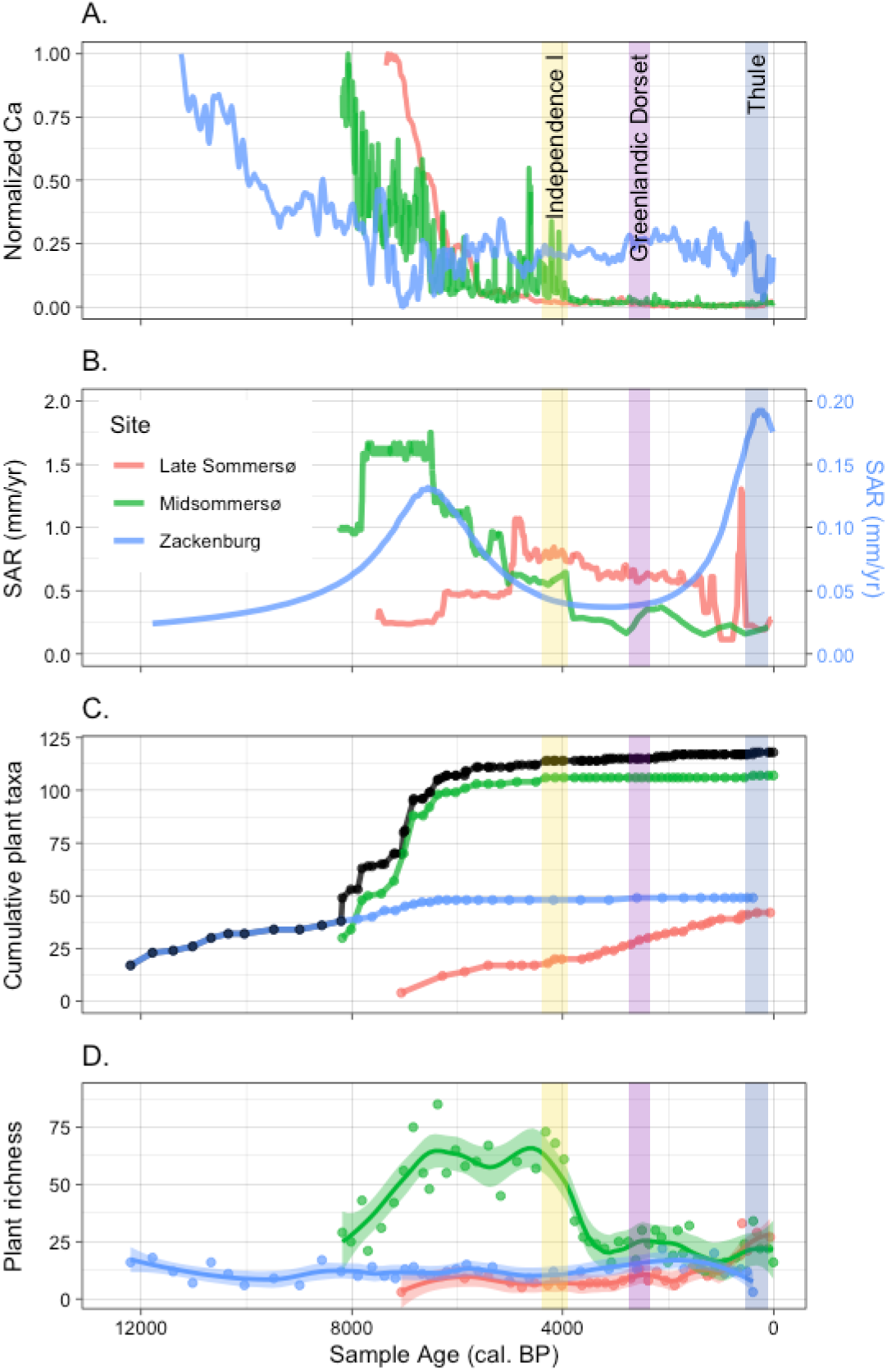
Temporal changes in plant richness and sediment properties over the study period at three lakes studied in northeast Greenland; (A) Changes in sediment properties measured as Ca normalised to 0-1, (B) sediment accumulation rate (SAR) expressed as mm/yr. Note: Zackenberg uses the second y-axis. (C) cumulative plant richness over time with the total from all three sites plotted in black, and (D) total number of plant taxa detected per sample (Hill q=0) from metabarcoding data. Cultural periods are indicated as colored vertical bars labelled in the panel (A).

### Plant and animal DNA

Metabarcoding recovered 121 native plant and nine native animal taxa across the three records, with no native taxa detected in any of the negative controls. Shotgun metagenomic sequencing of selected samples corroborated the dominant metabarcoding detections and showed DNA damage patterns consistent with the presence of authentic ancient DNA (Supplementary Information 5, Supplementary Table 16). There was high overlap in taxa detected between the two methods, and damage patterns were observed for all taxa frequently detected in the metabarcoding data. Only two forbs and fox were unique to the shotgun data. The metabarcoding dataset provides overall higher richness, taxonomic, and temporal resolution and thus was used for further downstream analysis (Supplementary Information 2-6, Supplementary Tables S6-S10).

The plant record includes algae, aquatic macrophytes, bryophytes, dwarf shrubs, ferns, forbs, graminoids, and horsetails. It provides earlier regional records for 15 taxa and the fifrst northeast Greenland paleoecological records for ∼40 taxa (Supplementary Information 3, Supplementary Table S11). We computed CONISS zones (Constrained Incremental Sum of Squares cluster analysis) and identified site-specific vegetation shifts from ∼8400 cal BP onward. Most shifts did not coincide with cultural periods, except for a major change at Midsommersø coinciding with the end of the Independence culture around 3900 cal BP (Supplementary Information 3, Supplementary Table S12).

About 90% of the plants were recorded prior to 6400 cal BP. Plant richness was roughly twice as high at Midsommersø as at the other lakes, peaking during the Mid Holocene and early Late Holocene before declining sharply around 3900 cal BP. In contrast, richness remained relatively stable at Zackenberg and increased only during the late Holocene at Late Sommersø (Fig. 2, Supplementary Information 3).

The mammal record was dominated by muskox (detected in 40 out of 50 samples at Midsommersø), Arctic hare (*Lepus arcticus*) and northern collared lemming (*Dicrostonyx groenlandicus*), with scattered records of reindeer in older samples. Two genera of geese were recorded; the more northern genus *Branta* being more frequent than *Anser.* We detected rock ptarmigan (*Lagopus muta*) in five samples from 6500-4000 cal BP at Wandel Dal. Arctic char (*Salvenius alpinus*) was detected at both lakes in Wandel Dal as well as a single sample from Zackenberg. We also detected one terrestrial and one aquatic worm (Fig. 3, Supplementary Data 1, Supplementary Information 4, Supplementary Table S13).

**Fig. 3.**
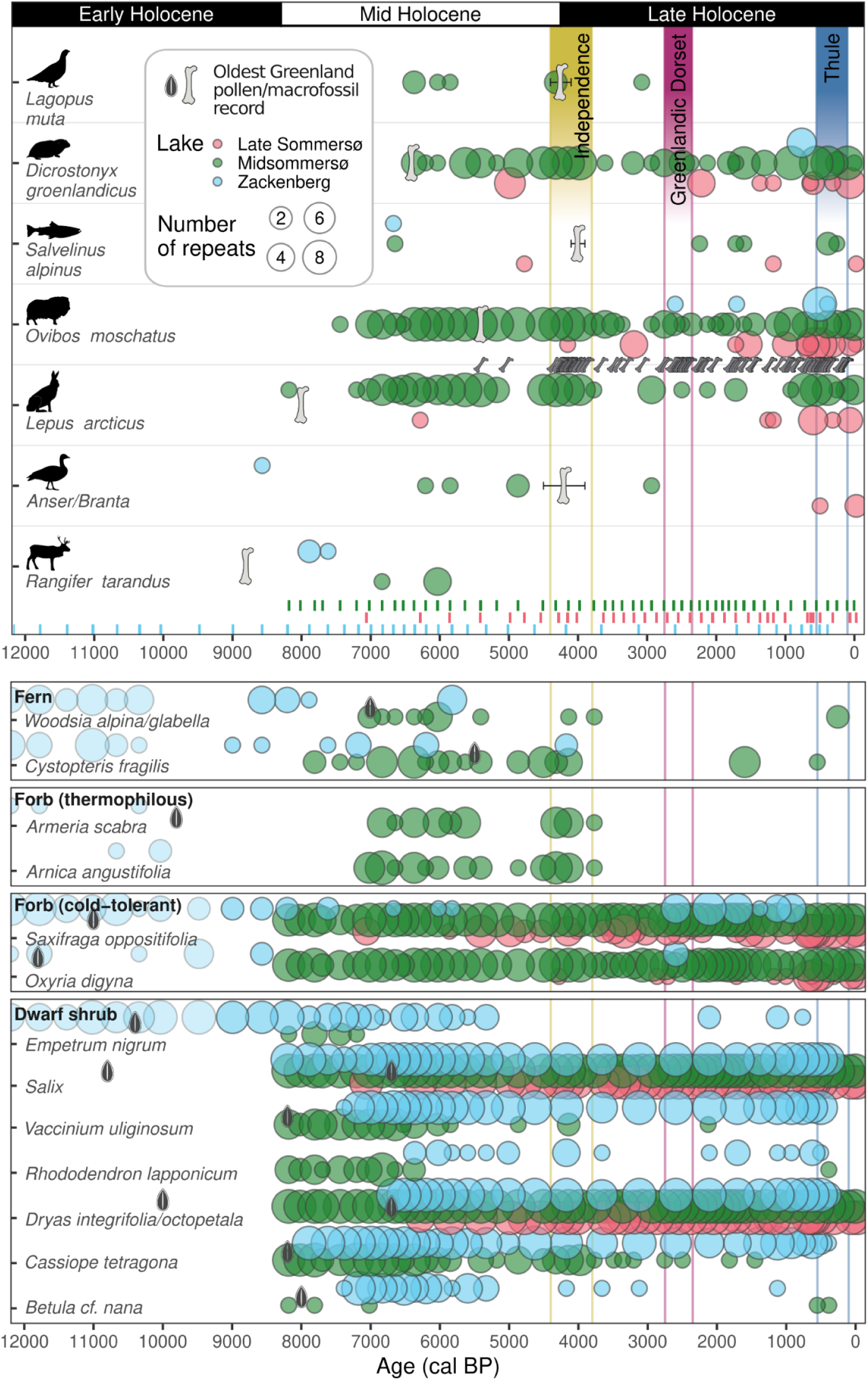
The plants and animals recorded in two lakes from Wandel Dal (Late Sommersø and Midsommersø) and one lake at Zackenberg, with the observed number of PCR repeats (1–8). Samples are indicated by vertical bars. The oldest post-glacial macrofossil and pollen records in Greenland are shown for comparison. For Salix and Dryas, two possible oldest pollen taxa within the group are represented. For muskox, all known macrofossil records in Greenland are presented as smaller markers (Supplementary Table S11, S13-14).

### Late Glacial and Early Holocene

Only the core from Zackenberg covers these periods. Twenty-three plant taxa appeared in the Late Glacial (12,200-11,700 cal BP; Figs. 2-3, Supplementary Information 3, Supplementary Table S11). These were mostly forbs, graminoids, and bryophytes that today are widespread in the circumpolar High-Arctic area, but also included two fern taxa and the relatively warmth-demanding forb, *Armeria scabra*. Moreover, we note a high abundance of the Low-Arctic, bird-dispersed, dwarf shrub *Empetrum nigrum* (Fig. 4), which was already found in the oldest sample dating to 12,200 cal BP (Fig. 3).

**Fig. 4.**
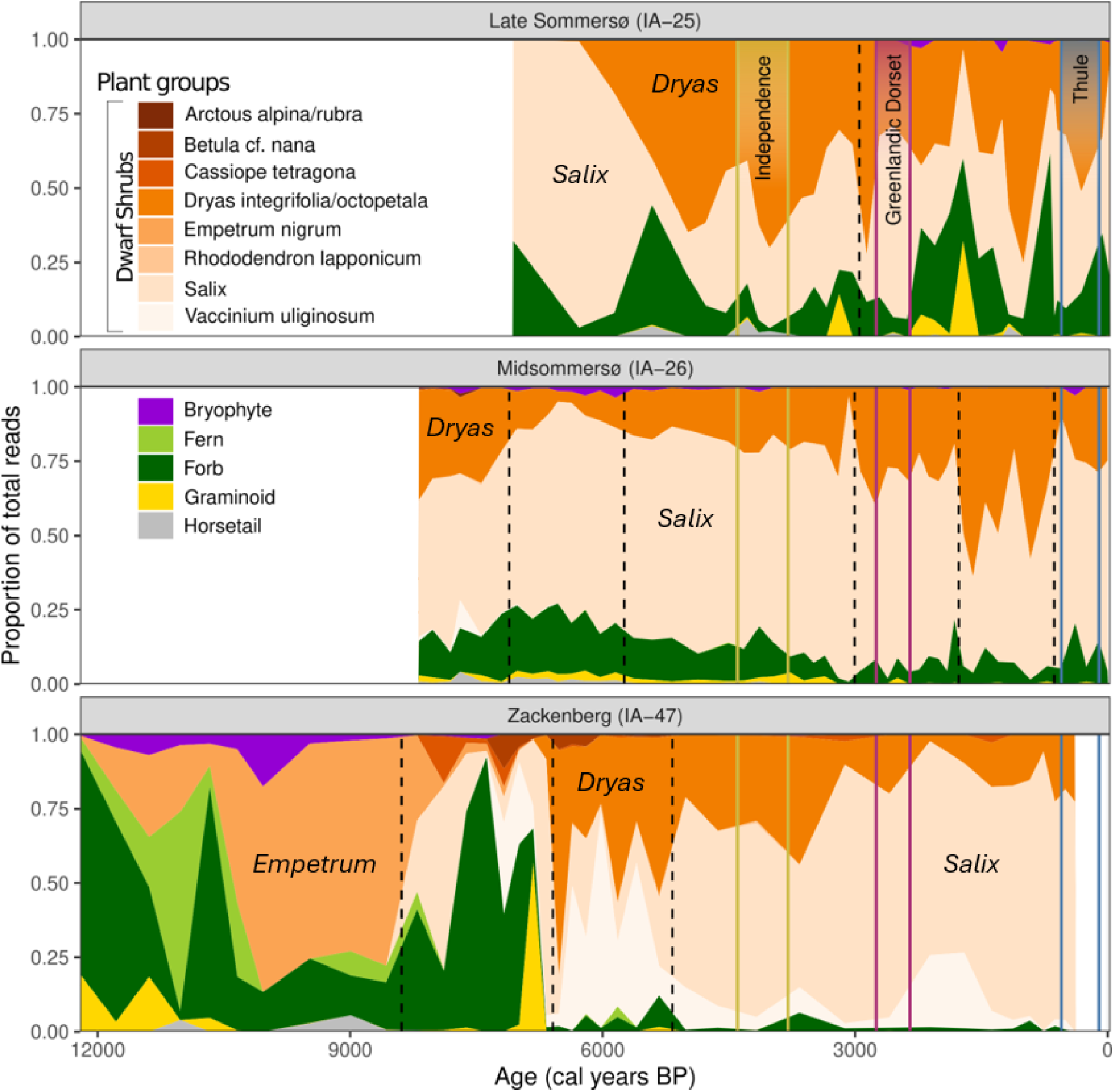
Changes in relative abundance of terrestrial plant growth forms over time estimated as the proportion of DNA reads for the three studied cores. CONISS breakpoints from the lowest taxonomic resolution of the data are indicated as vertical discontinuous lines. See also Supplementary Fig. 3.3 for the same plot using proportion of PCR repeats and thus emphasising species diversity.

A further 15 plant taxa appeared in the Early Holocene (11,700–8300 cal BP), including the thermally restricted forb, *Arnica angustifolia* (Figs. 2-3). No mammals were found in this early part of the sedaDNA record, but a reindeer antler found about 50 km to the east has been dated to 8800 cal BP. Geese (*Anser*) are found in one sample dated to 8600 cal BP (Fig. 2).

### Mid Holocene and Independence culture

The Zackenberg core spans the entirety of the Mid Holocene (8300–4250 cal BP) and Independence culture (4500–3900 cal BP) periods, whereas both Wandel Dal cores capture the entire Independence period and most of the Mid Holocene. The 78 newly detected plant taxa partly reflect the inclusion of the two Wandel Dal sites (Figs. 2 and 3, Supplementary Fig. 3.1). *Salix* appeared at both Zackenberg and Wandel Dal by 8200 cal BP, and caused a major turnover in dominant dwarf shrubs at the former (Fig. 4). At Wandel Dal, the characteristic dwarf shrubs of the northern and middle arctic tundra zones, *Dryas* and *Cassiope tetragona* respectively, together with the Low-Arctic dwarf shrubs *Vaccinium uliginosum, Rhododendron lapponicum, Betula* cf. *nana* and *Arctos alpina* were present from the earliest sample 8200 cal BP, whereas they appeared somewhat later at Zackenberg. Moreover, a range of Low-Arctic forbs such as *Pyrola grandiflora* and *Saxifraga aizoides*, together with the aquatic *Hippuris lanceolata* and the sedge *Juncus arcticus* were found up to 800 km north of their current distribution (Supplementary Fig. 3.4), indicating substantially warmer conditions than today. At Midsommersø, vegetation composition and relative abundance shifted markedly around 7100 cal BP, accompanied by the arrival of numerous bryophytes and graminoids and a peak richness of 83 taxa at 6800 cal BP (Figs. 2–4; Supplementary Information S3; Supplementary Table S11). This turnover included declines in several thermophilous dwarf shrubs and the appearance of both warmth-demanding and cold-tolerant forbs, indicating a substantial reorganisation of the vegetation rather than a clear transition to a more cold-tolerant flora.

Reindeer DNA was found in two samples from Zackenberg (7900 and 7600 cal BP) and two from Midsommersø (6800 and 6000 cal BP), thus somewhat younger than the oldest dated antler (Fig. 3, Supplementary Table S13). Muskox DNA was found in 17 of 18 samples from Midsommersø dated to 7400-3900 cal BP, and detected in all PCR repeats from 6400-4100 cal BP. It was also found scattered in Late Sommersø, but not at Zackenberg during the Mid Holocene. Arctic hare and northern collared lemming are found in most samples from Midsommersø, are scattered in Late Sommersø, but with only a single detection of the latter at Zackenberg. Geese and rock ptarmigan were found in five samples each from Midsommersø, whereas Arctic char was detected in one sample from each of the three lakes for this period. Our bird and fish detections predate fossil records by 2000-2600 years (Fig. 3).

The arrival of people of the Independence culture does not coincide with any major change in sedimentation rate or sediment properties, plant CONISS zones or the arrival of game animals. However, plant richness and game availability were both high at this time (Figs. 2-3). Some of the most thermophilic dwarf shrubs, such as *Empetrum nigrum, Rhododendron lapponicum* and *Betula* cf. *nana* had already disappeared, suggesting a climate cooler than the Holocene Thermal Optimum of the Early to Mid Holocene. *Eutrema edwardsii* (“Arctic wasabi”) was the only plant that appeared for the first time during the Independence occupation.

### The end of Independence cultural presence

At the end of the Independence culture (3900 cal BP), plant richness at Midsommersø abruptly decreased and there was a shift in floral composition (Fig. 2. Supplementary Figs. 3.1 and 3.3a,b). The transition is marked by a major sedimentological transition with possible linkage to lowered lake level at Midsommersø (Fig. 2. Supplementary Information 1). In contrast, plant richness remained stable and low at the two other sites (Fig. 2), with no change in species composition or relative abundance (Fig. 4, Supplementary Fig. 3.3). Muskox did occur in the subsequent samples at Midsommersø dated to 3976, 3771, 3610, 3492, and 3362 cal BP. However, rock ptarmigan and geese were only found in a single sample after 3900 cal BP, and Arctic hare was not detected in either of the Wandel Dal cores between 3800 and 2900 cal BP (Fig. 3, Supplementary Fig. 4.1).

### Greenlandic Dorset Cultural period

No specific changes were seen in the animal data at the start or end of Greenlandic Dorset culture 2800-2400 cal BP at Wandel Dal, but the start does coincide with the first detection of muskox at Zackenberg (Fig. 3). There was a significant change in the vegetation at Midsommersø and Late Sommersø ∼3000 cal BP (Fig. 4), ∼200 years before the start of the Greenlandic Dorset culture. The proportion of *Dryas* increases in both lakes, and *Salix* increases at Late Sommersø (Supplementary Fig. 3.2).

### Thule Cultural period

Significant vegetation changes occurred in both Wandel Dal lakes coinciding with the onset of the Thule cultural period around 700-600 cal BP (1400 CE). Plant richness increased, particularly at Late Sommersø, while Midsommersø showed a decline in *Salix*, an increase in *Dryas*, and the reappearance of several graminoids and forbs (Figs. 3 and 4). This period also coincided with more consistent detections of Arctic hare at Late Sommersø.

## Discussion

Our records reveal that muskox were established in northeast Greenland several millennia before the arrival of people of the Independence culture. They also indicate that the region supported a richer flora and fauna under warmer climatic conditions during much of the Early and Mid Holocene as compared to the Late Holocene.

### Post-glacial vegetation assembly and early warming

The Zackenberg record provides a unique opportunity to reconstruct the initial stages of postglacial vegetation, as it extends back to local deglaciation. In Wandel Dal deglaciation may predate the oldest recovered sediments by 1–2 millennia (24). The earliest plant assemblages at Zackenberg were dominated by widespread, High-Arctic bryophytes, forbs, graminoids, and ferns. This pattern of postglacial colonization is also supported by previous macrofossil and pollen studies from Greenland (2, 11, 12) and sedaDNA records from northern Fennoscandia (25). However, several detections at Zackenberg point to relatively favorable Late Glacial and Early Holocene conditions. Most notable, *Empetrum* rather warmth-demanding forbs and ferns occurred in the oldest rock-flour glacial deposits, dated to 12,200 cal BP, 1800 years before the region’s earliest dated macrofossil (12). Despite greater chronological uncertainties at the sequence basel, our result may indicate that warming started earlier than shown by other proxies (12, 26).

The establishment and persistence of *Salix* marks an important ecological development. *Salix* at Zackenberg recorded from 8200 cal BP most likely represents the Low-Arctic dwarf shrub *S. herbacea,* which has regional macrofossils extending to 10,800 cal BP (2). In Wandel Dal, it likely represents the High-Arctic dwarf shrub *S. arctica*, the only species currently north of 78°N in Greenland and previously recorded in nearby macrofossil and pollen sequences from c. 7800 cal BP (27–29). Its persistence throughout the Midsommersø record contrasts with its Late Holocene absence in coastal records (27, 29, 30). The establishment of *Salix* may have contributed to the development of terrestrial ecosystems capable of supporting resident herbivores, including muskox, reindeer, and ptarmigan, while also providing a potential fuel source for human populations (16, 31).

Regional differences in *Dryas* arrival may reflect dispersal history more than climatic suitability; in northern Fennoscandia and the Polar Urals, *Dryas* preceded most other dwarf shrubs (25, 32). The sedaDNA records in Wandel Dal may reflect immigration of the North American *D. integrifolia,* which currently is dominant in Wandel Dal, whereas the late arrival in Zackenberg may reflect chance dispersal across the Atlantic of the Eurasian *D. octopetala,* which is dominant at Zackenberg today. Macrofossils of *D. integrifolia* were found from approximately 6800-4100 cal BP in the Skallingen area south of Wandel Dal (29), whereas *D. octopetala* has been found from 7700 cal BP south of Zackenberg (33). Thus, long migration routes, rather than dispersal ability or climate, may have constrained its spread across the Arctic.

The occurrences in Wandel Dal of numerous Low-Arctic dwarf shrubs, forbs, sedges, and aquatic plants far north of their present distributions provide strong evidence for a warmer Early and Mid Holocene climate. This interpretation accords with previous regional evidence for warmer-than-present conditions, including records of *Armeria scabra* and *Salix arctica* beyond their modern distributions (12, 27, 34). The vegetation changes at Midsommersø around 7100 cal BP nevertheless indicate that climate was not the only control on local plant composition. The decline of several thermophilous dwarf shrubs alongside the appearance of many new taxa including both warm- and cold-adapted forbs suggests changing snow cover, soil moisture, catchment development, or herbivore pressure.

### Faunal assemblage and vegetation changes

The faunal record suggests that reindeer and geese were among the earliest vertebrates, followed by Arctic hare, muskox, northern collared lemming, and rock ptarmigan. Similar early occurrences of reindeer and geese have been reported from northern Fennoscandia (35). Reindeer were detected in samples dated 8000-6000 cal BP in both Wandel Dal and Zackenberg. Its absence from younger records parallels the scarcity of reindeer remains in archaeological assemblages (1, 12, 20), and suggests that reindeer were not an important or reliable resource during most periods of human occupation. This interpretation contrasts with the notion that the Independence Culture simply did not hunt reindeer. Instead, reindeer may have been an early but transient component of the fauna, whereas muskox became the dominant large terrestrial herbivore by the time the first humans arrived.

Muskox and hare dominated both Wandel Dal assemblages, consistent with an opportunistic hunting strategy (1). In contrast, Arctic fox was detected only once in the shotgun metagenomic data and was absent from metabarcoding records despite being common in Independence bone assemblages. This discrepancy is likely methodological as Arctic fox is also absent from other arctic sedaDNA records (10, 35). Conversely, frequent lemming detections, despite their absence from archaeological assemblages, suggest that these small rodents were locally abundant but not exploited by human groups.

The change in both plant species composition and abundance around 7000 cal BP coincided with the arrival of most of the regional vertebrates. Vertebrates may have promoted the dispersal of plants and thereby have contributed to higher plant richness (36). Also, the decrease in dwarf shrubs and increase in forbs may relate to grazing (37). This is in contrast to northern Fennoscandia, where the major change in vegetation was prior to herbivore arrival and where herbivores seem to have had a low impact on vegetation composition (25, 35). A possible explanation for this is the high density of muskox, as suggested by the high replication and detection in both metabarcoding and shotgun data. This consistent detection rate for a mammal is higher than any other Arctic studies that we are aware of (10, 35) and is commonly only observed in sedaDNA studies of domestic animals (38). Thus, we infer a high population density of muskox, which could have caused the observed change in the vegetation.

### Muskox persistence and human occupation

The continuous muskox record from Midsommersø, beginning 7400 cal BP, shows that muskox were established in northern Greenland long before Independence settlement, predating the earliest reported muskox bones from Peary Land by at least 2000 years and the main concentration of early archaeological remains by approximately 3000 years (1, 15, 17, 18). The apparent near-synchrony of muskox and human arrival inferred from archaeological bone assemblages therefore reflects the sampling bias at occupation sites rather than simultaneous colonization. Although muskox were an important resource for human entry into northeast Greenland, our findings do not support a model in which people directly followed muskox during their initial expansion into and across northern Greenland. The long interval between muskox establishment and Independence settlement indicates that prey availability alone did not determine the timing of human arrival; climate and sea-ice conditions, travel routes, technological capacity, demographic processes, and social connections with Arctic Canada likely also mattered.

Muskox detections became less consistent after approximately 3900 cal BP, but this pattern does not demonstrate local extinction. Mammalian sedaDNA is inherently difficult to recover (9) as evident from the lack of muskox detection in 535 samples from 74 circumarctic sites in, despite bone records suggesting its presence (10, 15). Further, 82% (87% from 7400 cal BP) of our samples from Midsommersø had muskox, whereas only 63% of samples from the circumarctic region had any mammals at all, reflecting the generally low probability of detecting mammals even when they are present (10). Some of the youngest radiocarbon dates for Independence I sites farther southeast may indicate gradual expansion accompanied by local hunting-related depletion. Nevertheless, although populations likely fluctuated, our data do not support muskox loss as the principal cause of the Independence decline.

The major ecological transition at approximately 3900 cal BP, particularly evident at Midsommersø in sedimentology (24) and vegetation, coincides with the end of Independence occupation. Reduced plant richness, altered vegetation, and lower sedimentation rates indicate environmental deterioration. Although this abrupt change was recorded in our Zackenberg record, cooling has also been inferred there 4500-3500 cal BP (12). Temporary loss or reduced detection of Arctic hare, geese, and ptarmigan suggest changes in smaller or seasonal resources. Environmental change may therefore have reduced resource predictability and diversity without eliminating muskox.

The first muskox detection at Zackenberg broadly coincides with Greenlandic Dorset occupation. A gap in muskox bone records between 76.28° N (Store Koldeway) and 71.5° N with only recent dates further south (Fig. 1), has been attributed to scarcity of archaeological sites as there are no obvious dispersal barriers that would prevent southward migration (18). However, our late detection of muskox DNA in Zackenberg supports delayed expansion rather than solely uneven archaeological sampling. Such a distributional limit could have influenced the southern extent of Independence settlement, although additional records are needed to establish this relationship. Because the lake at Zackenberg is situated at higher altitude and away from most Wollaston Forland archaeological finds (12) and it may not fully represent conditions encountered by early humans.

Finally, the Thule period (∼1400 CE) coincides with renewed changes in vegetation and increased mammal sedaDNA detections at Wandel Dal, suggesting increased ecological richness. However, Thule arrival was not necessarily driven by climate or terrestrial game availability: their distinct technologies and greater reliance on marine resources made them less dependent on terrestrial game than earlier groups.

## Conclusion

Our records reveal that muskox were established in northeast Greenland several millennia before the arrival of the first humans, challenging the Muskox Way hypothesis that Independence I populations followed migrating muskox herds into the region. During the Early and Mid Holocene, northeast Greenland supported diverse terrestrial plant communities, including species occurring hundreds of kilometres north of their present-day distributions. The end of the Independence occupation around 3900 cal BP coincided with a major ecological transition, yet muskox persisted, suggesting that their local disappearance was not responsible for the abandonment of the region.

## Methods

### Sites

#### Wandel Dal: Midsommersø and Late Sommersø

Midsommersø and Late Sommersø (unofficial name) are located within Wandel Dal in Inutoqqat Nunaat (Peary Land), in the High-Arctic of northern Greenland (Fig. 1). Wandel Dal connects J.P. Koch Fjord to the west with Jørgen Brønlund Fjord, a tributary of Independence Fjord, to the east. The valley also has an important human history and has been a corridor for human migration since c. 4500 cal BP (1, 3, 6, 16). Wandel Dal is within Bioclimate Subzone B, which is characterized by plant cover that is 5-25% vascular plants and up to 60% cryptogams (39). More specifically, vegetated areas of Subzone B are generally a mixture of rush/grass cryptogam tundra, prostrate dwarf shrub, herb and forb tundra (39).

Midsommersø (82° 15’ N, 34° 15’ W; 80 m a.s.l.) is a large lake system that has a 5,400 km^2^ catchment area and receives meltwater from sectors of the Greenland Ice Sheet, and the Hans Tausen, Bure, and Storm Ice Caps (Fig. 1). The lake consists of two basins: Øvre (upper) Midsommersø with a surface area of ∼58 km^2^, and Nedre (lower) Midsommersø with a surface area of ∼26 km^2^. Our record is from the central basin of Nedre Midsommersø with a water depth of 45 m (Fig. 1). Late Sommersø (82° 13.73’ N, 33° 31.512’ W; 120 m a.s.l.) is a small (0.05 km^2^ surface area), shallow (9 m), closed basin lake located along the southern edge of Midsommersø (Fig. 1, Supplementary Table S1, Supplementary Fig. S1.2).

#### Zackenberg

Zackenberg 06 Lake (unofficial name, 74° 28.909’ N, 21° 29.245’ W) is the headwater lake of the glacial Store Sødal valley, which runs parallel to Tyroler Fjord, in the northeast Greenland National Park (Kalaallit Nunaanni nuna eqqissisimatitaq). The lake is 14 m deep, 1,250 m wide, and 600 m long, and is situated at approximately 265 m a.s.l. (Fig. 1, Supplementary Table S1). Its name is unofficial and comes from its proximity to the Danish Zackenberg Research Station in the Zackenberg Valley.

The lake lies within bioclimatic subzone C, the Mid Arctic Tundra Zone (39). Most lakes in this region exhibit moss-covered bottoms, while others host vascular plants such as the *Hippuris vulgaris* and the High-Arctic *Ranunculus hyperboreus* in their littoral zones (40). Ice typically begins to melt along the shores in mid-June, but the lakes are generally ice-free only from mid-late July until mid-late September.

### Sediment cores and chronology

#### Wandel Dal: Midsommersø and Late Sommersø

Lake coring was done in August 2022 from a floating platform using a gravity corer to recover the water-sediment interface and a piston corer to recover the full post-glacial sequence. We constructed a composite core from each site based on XRF profile scans and visual stratigraphic changes (Supplementary Information S1).

At Late Sommersø, we collected one surface core (LSD-22-01) and a longer piston core (LSP-22-03) to construct a 383-cm composite sequence. An age-depth model was generated based on fifteen radiocarbon measurements using the Bacon age-modelling software in R (41) (Supplementary Information S1).

The sediment record from Midsommersø is a 559-cm composite sequence that spans the last c. 8200 cal BP (24). An age-depth model was generated using the Bacon age-modelling software in R (41) with results from the analysis of anthropogenic Cesium (^137^Cs) and thirteen radiocarbon measurements (Supplementary Information S1). Details of the stratigraphy and chronology are presented by Balascio et al. (2026), but the record is generally defined by three lithostratigraphic units marked by compositional and sedimentation rate changes around 6500 and 3900 cal BP.

#### Zackenberg

The lake was cored from lake ice in 2019 using a 6cm in diameter UWITEC gravity corer. The sediments of the 80-cm-long ZAC1906-02 core is assumed to represent the complete sedimentary infill of the lake. The age–depth model was constructed using seven radiocarbon dates using the R “clam” package (42). The very low content of palynological remains prevented the use of this fraction for dating, and therefore the bulk organic fraction was instead used.

### Sedimentary ancient DNA

To minimize the risk of false positives, we took necessary steps all along the process from laboratory work to the final stage of bioinformatic and manual cross-checking of the data following (35). We used two different DNA approaches to obtain a robust dataset. All samples were analysed with the high-throughput and cost-efficient DNA metabarcoding method, which has the advantage of providing the highest detectability and taxonomic resolution for plants (21). Authenticity of ancient DNA was evaluated through extensive replication and use of negative controls (35). We complemented this method with shotgun metagenomic sequencing of 32 samples to characterize ancient DNA damage patterns.

#### DNA sampling and extraction

We took 50, 38 and 40 samples from the Midsommersø, Late Sommersø, and Zackenberg Lake 06 cores respectively; the former two in a laminar flow hood in the Quaternary laboratory at the University of Bergen, and Zackenberg at the Geosciences Barcelona

Lacustrine Sedimentology laboratory, using exposed DNA-free water as controls for DNA contamination in the sampling facilities. DNA extraction, metabarcoding, and library preparation were done in dedicated ancient DNA laboratories at UiT - The Arctic University of Norway. Initially, 8-10 g of sediments were homogenised and DNA was extracted from subsamples of ∼0.3 g using the DNeasy PowerLyzer PowerSoil kit (Qiagen), incorporating a bead beating step as in (43).

#### DNA metabarcoding

DNA was amplified using plant and mammal primers. For plants, the g–h primer pair, which targeting the P6 loop of the chloroplast *trn*L (UAA) intron in vascular plants was used (44). For mammals, a section of the mitochondrial 16S region was amplified using the MamP007F primers, which can also amplify other vertebrates as bycatch (45, 46). In mammalian metabarcoding, human DNA originating from reagents or laboratory personnel may be co-amplified with target DNA during PCR, thereby reducing the representation of low abundant wildlife DNA in the final sequencing read. To reduce the amplification of human DNA, we applied a dual-blocking oligonucleotide strategy to inhibit human DNA amplification (47). Each DNA extract was amplified eight times to 1) improve confidence in detections, 2) increase the chance of detecting species represented by low DNA template quantities, and 3) allow the use of a more conservative PCR replicate-based quantification statistic than the proportion of reads (see pros and cons of each quantification methods discussed in (25)). Unique flanking identification tags were added to the 5’ end of the primers to enable pooling of multiple PCR reactions (Supplementary Table S3). PCR reaction and cycling conditions, amplicon cleanup, and library preparation followed (47). Sequencing was done on an Illumina NextSeq2000 for 2 x 150 paired-end cycles at the Genomics Support Center Tromsø, Faculty of Health Sciences, UiT – The Arctic University of Norway.

The sequence data were analysed with the OBITools software package (48) following the methods described in (47). Plant barcodes were compared against four DNA reference libraries: 1) PhyloNorway (25, 49), 2) ArcBorBryo, comprising 815 arctic (50) and 835 boreal (51) vascular plant taxa as well as 455 arctic-boreal bryophytes (52) from the circumpolar region (n=2280 sequences of which 1053 are unique), 3) PhyloAlps containing 4604 specimens of 4437 taxa collected in the Alps and Carpathians (47, 49) (https://data.phyloalps.org/browse/), and 4) EMBL (release 143, n=159,748 sequences of 74,936 taxa). The identified barcodes were filtered to only retain those that had a 100% match across the whole barcode to the reference library, a minimum of 3 reads per PCR replicate detection, and at least 10 reads and 1 replicate across the total dataset. Matches to PhyloNorway and/or ArcBorBryo were prioritised because these reference libraries provide good coverage of the Greenland flora. Sequences with a 100% match across the whole barcode to any of the four reference libraries were manually checked for consistent assignment across reference libraries and that the identified taxon was ecologically and biogeographically plausible. Sequences were then categorised as 1) plausible, 2) marginal, or 3) sequence probably representing common food contaminants, sequence error, and/or database error. Only sequences of category 1 are presented in the figures and text, whereas all three categories are given in the Supplementary Table S7. The quality of each sample was checked by plotting Metabarcoding Technical Quality scores (MTQ), Metabarcoding Analytical Quality scores (MAQ), total raw read count, and mean barcode sequence length following Rijal et al. (53).

The animal barcodes were identified with a reference library based on the EMBL nucleotide database collected on 2022-10-06, with additional 16S reference sequences added for *Lemmus lemmus* (from (46)), *L. sibiricus* and *Microtus oeconomus* (35). The identified barcodes were filtered to only retain those that had a 100% match across the whole barcode to the EMBL, with read and PCR repeat threshold as for plants.

The OBITools output was cross-checked using the Basic Local Alignment Search Tool (BLAST), with the blastn function. For sequences that had equal matches to several taxa, the taxa found in the region today were assumed to be the correct identification. When several sequences matched the same taxon, we checked if the less frequent variant mainly co-occurred with the more frequent variant, and, if so, they were assumed to be a sequence variant and so sequence counts were merged for quantitative analyses.

Wolf (*Canis lupus)* and dog (*Canis lupus familiaris*) share 16S barcodes. Sporadic occurrence of *Canis lupus* in four samples likely represents dog contamination, as this was also detected in one negative control. Further, one of these samples, IA-26-3_SED37, was also processed for shotgun metagenomic sequencing without detection of *Canis* and the only *Canis* detected in the shotgun analyses did not show an authentic ancient DNA damage pattern (Supplementary Fig. 5.3). *Canis* is therefore assumed to represent dog contamination.

#### DNA shotgun analyses

We shotgun sequenced 32 DNA extracts used for metabarcoding; three from Late Sommersø, 18 from Midsommersø, and 11 from Zackenberg. DNA extracts were transported from UiT to the Centre for Palaeogenetics in Stockholm where shotgun library preparation was conducted. Double-stranded libraries were prepared from 20 µL of each DNA extract following (54). Six negative sampling controls (four from Midsommersø and one from both Zackenberg and Late Sommersø) as well as a negative library control were processed alongside the 32 samples. The shotgun libraries were uniquely dual indexed and AMPure XP beads were used for purification and removal of suspected primer-dimers. The libraries were then pooled and sequenced on the Illumina NovaSeqX platform using 2 x 100 cycle chemistry at the Norwegian Sequencing Center, University of Oslo, yielding a total of 15.4 billion reads.

The paired-end read files had adapters removed, were merged, and had duplicate sequences removed using fastp (55). SGA preprocess was used to remove low complexity reads (dust filter = 4, (56). The filtered reads were multi-mapped to three different databases using bowtie2 (57); the bacterial genome taxonomy database (GTDB, release 232), the 1,541 assembled genome skims of arctic vascular plants from PhyloNorway (10), and a custom database of 42 metazoan genomes retrieved from EMBL of probable fauna and contaminants (Supplementary Table 15). Any mappings with ≥97% identity were retained and reads were assigned to the lowest common ancestor shared among all mappings using a custom python script. Taxonomic assignments were collapsed to the genus and family-levels with a minimum of 200 reads per sample. Bacterial assignments, as well as reads assigned at higher than family-level, were not considered for downstream analysis. As one safeguard to ensure reads were not only mapping to small portions of the reference genomes, a taxonomic assignment was only retained if >60% of the k-mers (k=31) of the assigned reads were unique which was calculated through a custom python script.

To support the ancient origin of DNA assigned to our taxa of interest, cytosine deamination patterns were characterized using pyDamage (58) grouping all reference contigs together (-g option). A taxonomic assignment was considered ancient if its assigned reads fit the GLM model of pyDamage with a p-value ≤ 0.05, a residual standard mean error ≤ 0.01, and a magnitude of damage > 0.02. The magnitude of damage is defined as the following where *CT*_0_ denotes proportion of C-T transitions at position 0 of the 5’ end of the fragment and *GA*_0_ denotes proportion of G-A transitions at position 0 of the 3’ end:

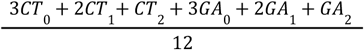

### Plant traits and distribution

Circumpolar and N Greenland vascular plant distribution as well as climate distribution types were taken from Bay (28). Global distributions for bryophytes were taken from GBIF if the metabarcode sequences were determined to the species level.

### Compilation of pollen, and plant and animal macrofossil records

Dates of first arrival for mammals, fish and birds were compiled from Bennike and Wagner (7). Additional dates on muskox were compiled from (1, 3, 15, 17, 18). Sampling coordinates for dates were compiled from the references above and by contacting the source museums cited in (15) - some of the latter have however been lost (Campos, pers comm. 2026) and are therefore not plotted. Compilations of first arrivals of plants were based on (59) with additions and cross-checking of further records (Supplementary Table S11). Grey literature was not used for this compilation.

## Supplementary Information

The supplementary information includes the following files:

Supplementary Information

Supplementary Tables 1-16

Supplementary Data 1 Metabarcoding

Supplementary Data 2 Shotgun metagenomics

## Data and software availability

The raw Illumina shotgun metagenomic and metabarcoding datasets are available from the European Nucleotide Archive (ENA) under accession PRJEB128621. Scripts used for processing metagenomic data can accessed at https://github.com/salanova-elliott/muskox_way

## Acknowledgement

We thank Frederik Fuuja Larsen and Christian Koch Madsen from Grøndlands Nationalmuseum for participation in fieldwork. We also thank Bjrøn Hattend and Linnea Betz, UiT, for help with figures. This research was funded by The European Research Council (ERC) under the European Union’s Horizon 2020 research and innovation programme grant agreement No 819192 for the IceAGenT project (IGA, MB, LD); grant TIME-LINES ERC-CoG-2021 101045309 (SN, IGA, SP, SG); and the UiT Arctic Ecosystem Genomic Centre (IGA, YL); the Zackenberg coring campaign and multiproxy characterization of the retrieved core were funded by the Spanish research projects PALEOGREEN (CTM2017-87976-P), NEOCLIM (PID2020-113798GB-C33), and HIGHCLIM (PID2024-163026OB-I00); the Wandel Dal campaign was jointly funded by the U.S. National Science Foundation (NSF) and the U.K. Natural Environment Research Council (NERC), including NSF grants OPP-2126047 (NLB), OPP-2125994 (RSB), OPP-2126212 (WJD), and NERC grant NE/W007541/1 (BP). Support was also provided by a U.S.-Norway Fulbright Scholar Award (NLB). LD, PDH and JA acknowledge support from the Knut and Alice Wallenberg Foundation (2022.0033).

## References

1. B. Grønnow, J. F. Jensen, The northernmost ruins of the globe. Eigil Knuth’s archeological investigations in Peary Land and adjajent areas of high arctic Greenland. Meddelelser om Grønland. Man and Society 29, 1–403 (2003).

2. O. Bennike, Colonisation of Greenland by plants and animals after the last ice age: a review. Polar Rec. 35, 323–336 (1999).

3. J. F. Jensen, A. B. Gotfredsen, First people and muskox hunting in northernmost Greenland. Acta Boreal. 39, 24–52 (2022).

4. H. P. Steensby, Contributions to the Ethnology and Anthropogeography of the Polar Eskimos. Meddelelser om Grønland 34 (1910).

5. H. P. Steensby, Origin of the Eskimo culture (Bianco Lunos Bogtrykkeri, 1916).

6. J. F. Jensen, U. Odgaard, S. Funder, P. Plumet, “First people in Greenland” in Human Colonization of the Arctic: The Interaction Between Early Migration and the Paleoenvironment, {V M. Kotlyakov, A. A. Velichko, S. A. Vasil’ev}, Eds. (Academic Press Inc. (London), 2017).

7. O. Bennike, B. Wagner, “Quaternary vertebrate from the North Atlantic islands” in Biogeography in the Sub-Arctic: The Past and Future of North Atlantic Biota, E. Panagiotakapulu, J. P. Sadler, Eds. (Wiley and Sons, 2021), pp. 147–160.

8. S. Garcés-Pastor, et al., “Terrestrial Plant DNA from Lake Sediments: Volume 6: Sedimentary DNA” in Tracking Environmental Change Using Lake Sediments, Developments in Paleoenvironmental Research., E. Capo, C. Barouillet, J. P. Smol, Eds. (Springer International Publishing, 2023), pp. 275–298.

9. T. J. Murchie, C. Giguet-Covex, P. D. Heintzman, V. Slon, Y. Wang, “Terrestrial fauna and hominin DNA from sedimentary archives” in Developments in Paleoenvironmental Research, (Springer International Publishing, 2023), pp. 299–378.

10. Y. Wang, et al., Late Quaternary dynamics of Arctic biota from ancient environmental genomics. Nature (2021). 10.1038/s41586-021-04016-x.

11. O. Bennike, et al., Early holocene plant and animal remains from North-east Greenland. J. Biogeogr. 26, 667–677 (1999).

12. O. Bennike, et al., “Late Quaternary Environmental and Cultural Changes in the Wollaston Forland Region, Northeast Greenland” in Advances in Ecological Research, T. R. C. B. E. M. C. F. Hans Meltofte, R. Morten, Eds. (Academic Press, 2008), pp. 45–79.

13. L. S. Epp, et al., Lake sediment multi-taxon DNA from North Greenland records early post-glacial appearance of vascular plants and accurately tracks environmental changes. Quat. Sci. Rev. 117, 152–163 (2015).

14. M. Sørensen, H. C. Gulløv, The prehistory of Inuit in northeast Greenland. Arctic Anthropol. 49, 88–104 (2012).

15. P. F. Campos, et al., Ancient DNA analyses exclude humans as the driving force behind late Pleistocene musk ox (Ovibos moschatus) population dynamics. Proc. Natl. Acad. Sci. U. S. A. 107, 5675–5680 (2010).

16. E. Knuth, Archeology of the musk-ox way. Contributions du Centre d’Études Arctique et Finno-Scandinaves (1967).

17. O. Bennike, Late Quaternary history of Washington Land, North Greenland. Boreas 31, 260–272 (2002).

18. O. Bennike, C. Andreasen, Radiocarbon dating of musk-ox (Ovibos moschatus) remains from northeast Greenland. Polar Rec. (Gr. Brit.) 41, 305–310 (2005).

19. M. Meldgaard, O. Bennike, Interglacial remains of caribou (*Rangifer tarandus*) and lemming (*Dicrostonyx torquatus*(?)) from North Greenland. Boreas 18, 359–366 (1989).

20. M. Meldgaard, The greenland caribou - zoogeography, taxonomy, and population dynamics. Meddr Grønland. Biosci 20 (1986).

21. N. A. Strandberg, et al., Capture probe, metabarcoding, or shotgun sequencing: Which best reflects local vegetation? Environ. DNA 8, e70270 (2026).

22. T. Leger, et al., Insights into the LGM-to-present evolution of the Greenland Ice Sheet from a data evaluated ensemble of numerical model simulations. (2024).

23. A. S. Dalton, et al., Deglaciation of the north American ice sheet complex in calendar years based on a comprehensive database of chronological data: NADI-1. Quat. Sci. Rev. 321, 108345 (2023).

24. N. L. Balascio, et al., Midsommersø records the Holocene glacial history of Wandel Dal, Inutoqqat Nunaat (Peary Land) northern Greenland. Quat. Sci. Rev. 378, 109874 (2026).

25. I. G. Alsos, et al., Postglacial species arrival and diversity buildup of northern ecosystems took millennia. Science Advances 8, eabo7434 (2022).

26. K. K. Andersen, et al., High-resolution record of Northern Hemisphere climate extending into the last interglacial period. Nature 431, 147–151 (2004).

27. B. Wagner, O. Bennike, Holocene history of Fiskesø, Prinsesse Ingeborg Halvø, eastern North Greenland. Bulletin of the Geological Society of Denmark 73, 209–220 (2024).

28. C. Bay, A phytogeographical study of vascular plants of northern Greenland - north of 74° northern latitude. *Meddelelser om Grønland*, Bioscience 36, 1–102 (1992).

29. S. Kusch, et al., Holocene environmental history in high-Arctic North Greenland revealed by a combined biomarker and macrofossil approach. Boreas 48, 273–286 (2019).

30. B. Wagner, O. Bennike, Holocene environmental change in the Skallingen area, eastern North Greenland, based on a lacustrine record. Boreas 44, 45–59 (2015).

31. D. R. Klein, C. Bay, Foraging dynamics of muskoxen in Peary Land, northern Greenland. Ecography (Cop.) 13, 269–280 (1990).

32. L. Topstad, et al., Vegetation history of the Polar Urals over the past 24,000 years based on sedaDNA from two well-dated lake archives. 386, 110024 (2026).

33. B. Wagner, O. L. E. Bennike, H. Cremer, M. Klug, Late Quaternary history of the Kap Mackenzie area, northeast Greenland. Boreas 39, 492–504 (2010).

34. B. Fredskild, Palynology and sediment slumping in a high arctic Greenland lake. Boreas 24, 345–354 (1995).

35. I. Alsos, et al., Biotic interactions may be more important than climate for shaping animal niches. Research Square (2026).

36. H. H. Bruun, R. Lundgren, M. Philipp, Enhancement of local species richness in tundra by seed dispersal through guts of muskox and barnacle goose. Oecologia 155, 101–110 (2008).

37. E. Post, et al., Large herbivores link plant phenology and abundance in Arctic tundra. PNAS Nexus 3, pgae454 (2024).

38. S. Garcés-Pastor, et al., Wild and domesticated animal abundance is associated with greater late-Holocene alpine plant diversity. Nat. Commun. 16, 3924 (2025).

39. D. A. Walker, et al., The circumpolar arctic vegetation map. J. Veg. Sci. 16, 267–282 (2005).

40. K. S. Christoffersen, S. L. Amsinck, F. Landkildehus, T. L. Lauridsen, E. Jeppesen, “Lake flora and fauna in relation to ice-melt, water temperature and chemistry at zackenberg” in *Advances in Ecological Research*, Advances in Ecological Research., (Elsevier, 2008), pp. 371–389.

41. M. Blaauw, J. A. Christen, Flexible paleoclimate age-depth models using an autoregressive gamma process. Bayesian Anal. 6, 457–474 (2011).

42. M. Blaauw, Methods and code for “classical” age-modelling of radiocarbon sequences. Quat. Geochronol. 5, 512–518 (2010).

43. I. G. Alsos, et al., Last Glacial Maximum environmental conditions at Andøya, northern Norway; evidence for a northern ice-edge ecological “hotspot.” Quat. Sci. Rev. 239, 106364 (2020).

44. P. Taberlet, et al., Power and limitations of the chloroplast trnL (UAA) intron for plant DNA barcoding. Nucleic Acids Res. 35, e14 (2007).

45. C. Giguet-Covex, et al., Long livestock farming history and human landscape shaping revealed by lake sediment DNA. Nat. Commun. 5 (2014).

46. T. Brown, et al., Paleoeconomy more than demography determined prehistoric human impact in Arctic Norway. PNAS Nexus 1, gac209 (2022).

47. S. Garcés-Pastor, et al., High resolution ancient sedimentary DNA shows that alpine plant diversity is associated with human land use and climate change. Nat. Commun. 13, 1–16 (2022).

48. F. Boyer, et al., OBITOOLS: a unix-inspired software package for DNA metabarcoding. Mol. Ecol. Resour. 16, 176–182 (2016).

49. I. G. Alsos, et al., The treasure vault can be opened: Large-scale genome skimming works well using herbarium and silica gel dried material. Plants 9, 432 (2020).

50. J. H. Sønstebø, et al., Using next-generation sequencing for molecular reconstruction of past Arctic vegetation and climate. Mol. Ecol. Resour. 10, 1009–1018 (2010).

51. E. Willerslev, et al., Fifty thousand years of Arctic vegetation and megafaunal diet. Nature 506, 47–51 (2014).

52. E. M. Soininen, et al., Highly overlapping diet in two sympatric lemming species during winter revealed by DNA metabarcoding. PLoS One 10, e0115335 (2015).

53. D. P. Rijal, et al., Sedimentary ancient DNA shows terrestrial plant richness continuously increased over the Holocene in northern Fennoscandia. Science Advances 7, eabf9557 (2021).

54. M. Meyer, M. Kircher, Illumina sequencing library preparation for highly multiplexed target capture and sequencing. Cold Spring Harb. Protoc. 2010, db.prot5448 (2010).

55. S. Chen, fastp 1.0: An ultra-fast all-round tool for FASTQ data quality control and preprocessing. Imeta 4, e70078 (2025).

56. J. T. Simpson, R. Durbin, Efficient de novo assembly of large genomes using compressed data structures. Genome Res. 22, 549–556 (2012).

57. B. Langmead, C. Wilks, V. Antonescu, R. Charles, Scaling read aligners to hundreds of threads on general-purpose processors. Bioinformatics 35, 421–432 (2019).

58. M. Borry, A. Hübner, A. B. Rohrlach, C. Warinner, PyDamage: automated ancient damage identification and estimation for contigs in ancient DNA de novo assembly. PeerJ 9, e11845 (2021).

59. I. G. Alsos, et al., The role of sea ice for vascular plant dispersal in the Arctic. Biol. Lett. 12 (2016).

