## Supplementary Information for "The Muskox Way: Humans, plants and animals in northeast Greenland through the Holocene"

#### 1 Age depth models and lithology

The age-depth models for Late Sommersø, Nedre Midsommersø and Zackeberg are based on 15, 13 and 7 radiocarbon measurements, respectively. For Midsommersø, anthropogenic Cesium ($^{137}Cs$) was also used and all dates for that site are presented in (1). For the other two sites, radiocarbon measurements are presented in Supplementary Table S2.

The 383 cm composite core from Late Sommersø consists of dense, minerogenic sediment at the base that increases in organic content. From 383 to 318 cm organic carbon values increase from c. 3% to 18%. The upper 318 cm are generally organic-rich with organic carbon values of c. 22% and include aquatic moss fragments throughout (Fig. S1.1). The age-depth model, developed using the Bacon software in R (2) and based on 15 radiocarbon dates, indicates low sedimentation rate (<0.05 cm/year) from ca. 7500-5000 cal BP, rising to a peak of ~0.10 cm/yr ca. 4900, possibly indicating a transition to greater runoff or higher autochthonous productivity or preservation. However, the increase in sedimentation rate ca. 5000 cal BP coincides with a drop in XRF-based Ca counts, suggesting either a decrease in photosynthetically driven precipitation of $CaCO_3$, reduced inwash of Ca-bearing minerogenic material to the lake from its watershed, or dilution by enhanced organic productivity at this time (Fig. 2, Supplementary Tables S1-4).

The sediment record from Nedre Midsommersø is a 559-cm composite sequence that spans the last c. 8200 years (Supplementary Fig. S1.1) Details of the stratigraphy and chronology are presented by Balascio et al. (2026). The age model from Nedre Midsommersø shows an overall pattern of increasing age with depth with the exception of two samples that are older than the surrounding sediment. We cannot rule out the potential influence of a hardwater effect since some of the plant material was unable to be identified, however the age reversals most likely reflect reworking of old plant material from the catchment. The record is generally defined by three lithostratigraphic units marked by compositional and sedimentation rate changes around 6500 and 3900 cal BP (Fig. 2, Supplementary Figs. S1.1). While the XRF-based Ca counts show similar patterns to the nearby Late Sommersø, the sedimentation rate is highest (>0.10 cm/year) 8200-5700 cal BP, decreases to <0.10 cm/year ca, 5700 cal BP, and to <0.05 ca. 3900 cal BP (Fig. 2).

The Zackenberg sediment record is 77.7 cm long and it spans the last ca. 12,000 years. This core can be subdivided, from bottom to top, into four lithological units. Unit 1, from the base of the core up to 75 cm of core depth, consists of light grey clays devoid of organic remains and corresponds to the initial rock-flour glacial deposits. Units 2 (75 - 28 cm) and 3 (28 - 2 cm) are intermediate organic-rich layers of light and dark brownish silts and clays: Unit 2 is finely laminated at millimetre scale, whereas Unit 3 is more distinctly banded at centimetre scale. The uppermost unit (Unit 4) comprises the top 2 cm of the core and consists of unconsolidated reddish silts and clays. An age–depth model, derived from seven radiocarbon dates and constructed using the latest version of the R ‘clam’ library (3), indicates that the record covers the entire Holocene, while its basal sediments extend into the Late Glacial (Supplementary Table S2, Fig. S1.1). The transition from glacial rock-flour sediments to the first organic-rich brownish layers dates to ca. 11,600 cal yr BP. Average sedimentation was about 0.0067 cm/year, increasing during 7000–5000 cal yr BP and the past 1500 years (Fig. 2, Supplementary Tables S1-S3, S5).

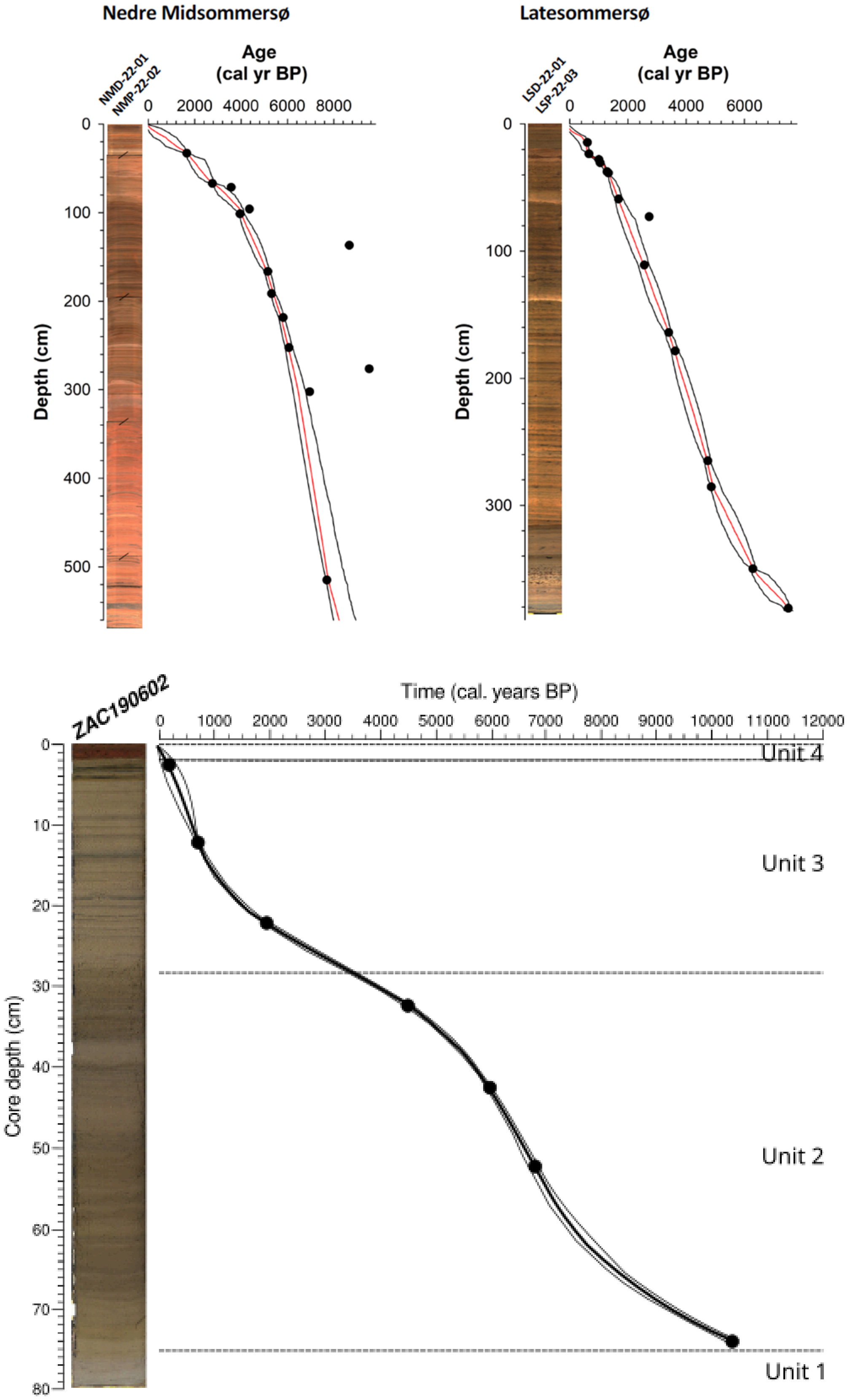


*Supplementary Fig. S1.1. Core images and age-depth models for Nedre Midsommersø, Late Sommersø and Zackenberg Lake 06. Dates disregarded in the age-depth model are shown as dots outside the 95% confidence interval.*

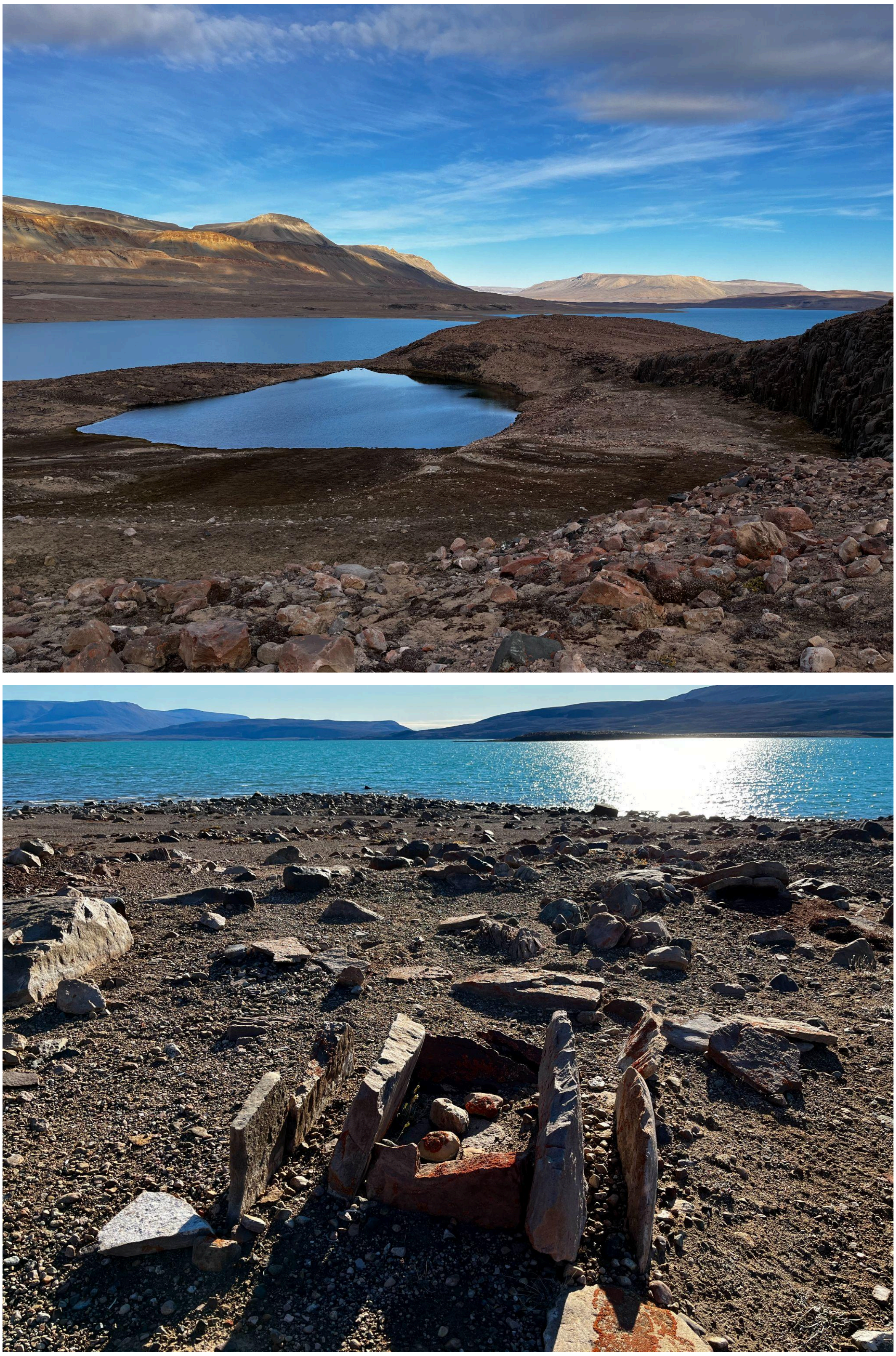

*Supplementary Fig. S1.2. The study sites in Wandel Dal, Inutoqqat Nunaat (Peary Land). Top photo: Late Sommersø in foreground and Nedre Midsommersø in background. Lower photo: An Independence hearth in front of Nedre Midsommersøfacing East from the terrace overlooking the bay. Photo credit: Bianca Perren.*

#### 2 Metabarcoding data and quality control

Our DNA metabarcoding of 128 samples and 48 controls resulted in a total of 53,453,330 and 27,633,529 raw reads for plants and animals, respectively. After final filtering, we retained 25 million plant reads and 2.1 million animal reads matching 121 native plants and 9 native animal taxa (Supplementary Data 1, Supplementary Tables S6-S12). No native plants or animals were detected in any negative controls (n=48 x 8 repeats = 384).

To explore whether any drop in species richness could be explained by the DNA quality of the samples, we plotted diagnostic plots using the P6loop data following Rijal et al. (4). They showed that the DNA quality was generally good for all three lakes. All except three samples from Late Sommersø had Metabarcoding Technical Quality scores (MTQ, reproducibility of the ten most read-abundant sequences) >0.5 and Metabarcoding Analytical Quality scores (MAQ, reproducibility of the ten most read-abundant sequences that are taxonomically identified) >0.18. All samples had MTQ and MAQ scores well above the negative controls that had MTQ and MAQ of <0.125 and <0.09, respectively. The exception were:

- IA-25-3_SED26_240213_LD_11 (no reads after OBItools in either P6loop or 16S)
- IA-25-3_SED27_240213_LD_12 (high number of *Nanochloropsis* algal reads in three repeats and one sequence identified to citrus in one repeat, zero reads in 16S after OBItools)
- IA-25-3_SED31_240213_LD_16 (five barcodes; three Cucurbita, 1 Fabaceae and 1 Solanoideae, so all assumed background contamination; 5 reads of human after Obitools).

These three samples were excluded from all further analyses.

In Nedre Midsommersø, the technical quality score is high throughout the period (Supplementary Fig. 2.1b). The Analytical Quality Score dropped about 3900 years ago (Supplementary Fig. 2.1d), coinciding with a species richness decreasing to around 25 taxa (Fig. 3). However, as this drop coincides with a drop in sedimentation rate, we interpret this as a change in climate rather than DNA quality. Change in sedimentation could cause changes in the catchment contributing to the DNA signal and in theory change the vegetation detected. However, we find it unlikely that so many species, including ubiquitous species, were clustered in a part of the catchment that had reduced contribution to the sediments after 3900 cal BP.

For Late Midtsommersø and Zackenberg, the MTQ and MAQ are somewhat lower than the Nedre Midtsommersø. As these lakes have smaller catchments and low species diversity (around 12 taxa), the likelihood of having 10 sequences well replicated per sample is low. Both these lakes have periods of shorter barcodes. However, these correspond to samples with a higher proportion of algae (Supplementary FIg. 3.3) and so the shorter barcodes reflect taxonomical composition rather than DNA fragmentation. The reproducibility is as can be expected for lakes with small flora, thus we regard the DNA quality as good.

The average time resolution was 202, 164, and 305 years for Late Sommersø, Midsommersø, and Zackenberg, respectively. When the two lakes from Wandel Dal were combined, the average time resolution was 96 years.

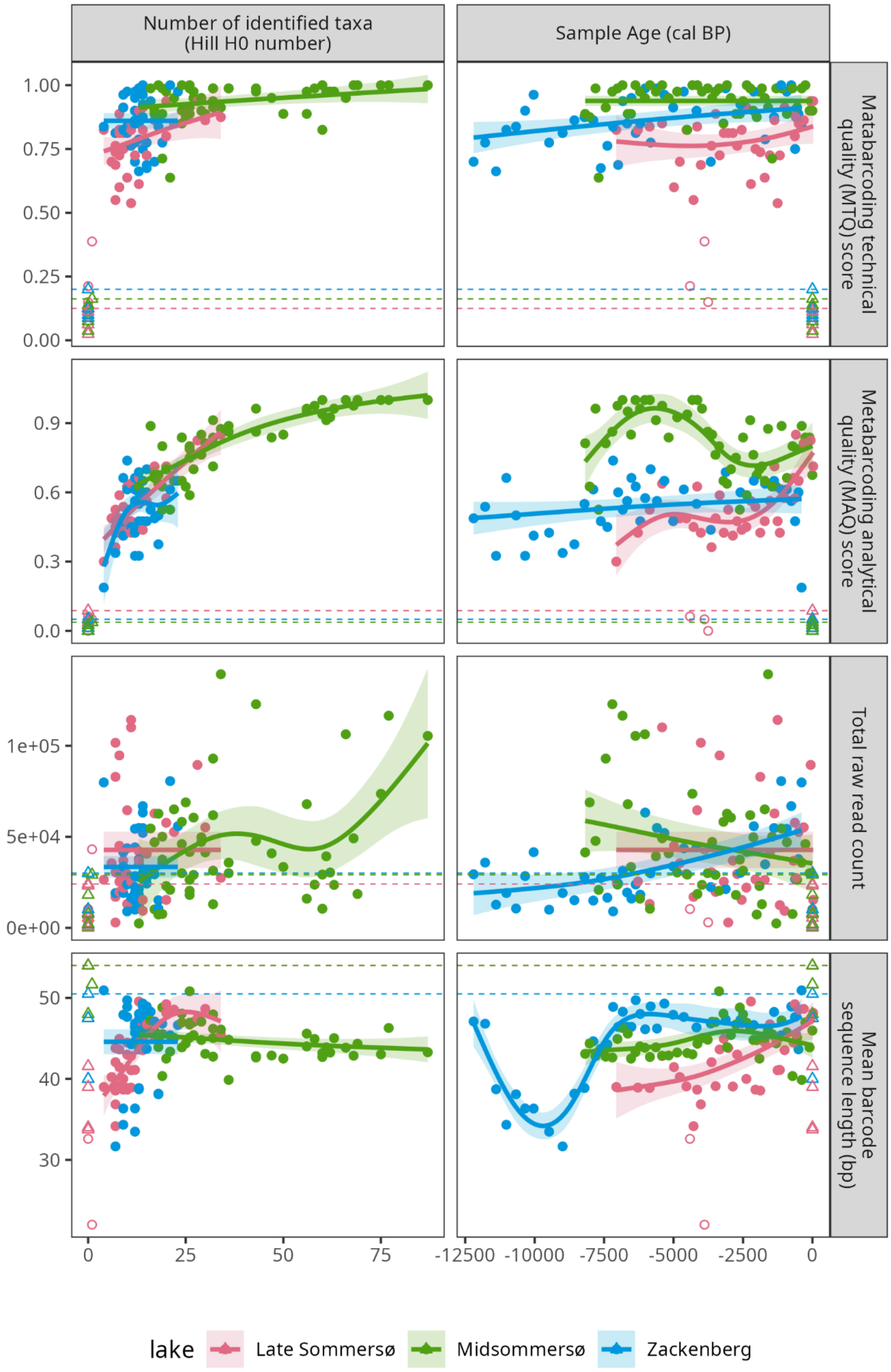


*Supplementary Fig. 2.1. Correlations between taxonomic richness (left) and time (right) against four measures of sedaDNA data quality. From top: 1) Metabarcoding Technical Quality score (MTQ), 2) Metabarcoding Analytical Quality score (MAQ), 3) total raw read count, and 4) mean barcode sequence length, across the entire dataset.Controls plotted at the zero line. Open circles represent samples with MTQ values < 0.5. These values are excluded from trendlines (GAM modelled). Triangles represent negative controls (plotted at 0 time). Dotted Lines represent the largest values found for controls.*

#### 3 The full plant metabarcoding record

We kept all 50 and 40 samples, respectively from each of Nedre Midtsommersø and Zackeberg and 37 of 40 from Late Midtsommersø. The plant metabarcoding record consisted of 4 taxa of algae, one aquatic macrophyte, 39 bryophytes, 8 dwarf shrubs, 2 ferns, 43 forbs, 22 graminoids and 2 horsetails.

The plant richness per sample was stable over 12,000 years at Zackenberg with an average of 12.1±3.7. At Late Sommersø, the average (excluding 3 poor samples) was 8.6±3.0 until about 1000 cal BP, after which it rose to 22.9±7.2. In contrast, Midsommersø had high richness in the Early and Mid Holocene (53.6±16.7) and this dropped steeply around 3900 cal BP to 22.8±6.2 (Fig. 2).

We compared our plant record with previous pollen and macrofossils although our compilation of plant records was less thorough than for mammals. Also, in contrast to mammal records where dating is often based on bones, the dates of first arrival of plants is commonly not given and has to be interpreted from diagrams. Further, differences in taxonomical resolution makes direct comparison difficult. Nevertheless, about half of the 78 vascular plants that we detected with metabarcoding have previously been recorded in palaeorecords from northeast Greenland. We, however, provide earlier appearance dates for 15 of them, for example *Empetrum nigrum, Bistorta vivipara*, and *Saxifraga oppositifolia.* In addition, we provide the first fossil record from northeast Greenland for ~40 taxa including arctic forbs such as *Eutrema edwardsii,* graminoids such as *Pleuropogon sabinei* and *Kobresia myosoides*, ferns such as *Cystopteris fragilis,* and dwarf shrubs such as *Rhododendron lapponicum* (Supplementary Table S11).

To find the main changes in the ecosystem, we computed CONISS zones based on the number of PCR repeats (thus giving weight to flora composition) and based on DNA reads (thus giving weight to abundance of dominant taxa) of the plants. Changes occurred at different times at Zackenberg, Midsommersø, and Late Sommersø (Supplementary Table S12, Supplementary Figs. 3.1-3.3). A major compositional shift at Midsommersø around 3,900 cal BP coincided with the end of Independence I, and changes in dominant taxa preceded the Dorset and Thule periods by roughly 100–200 years.

a)

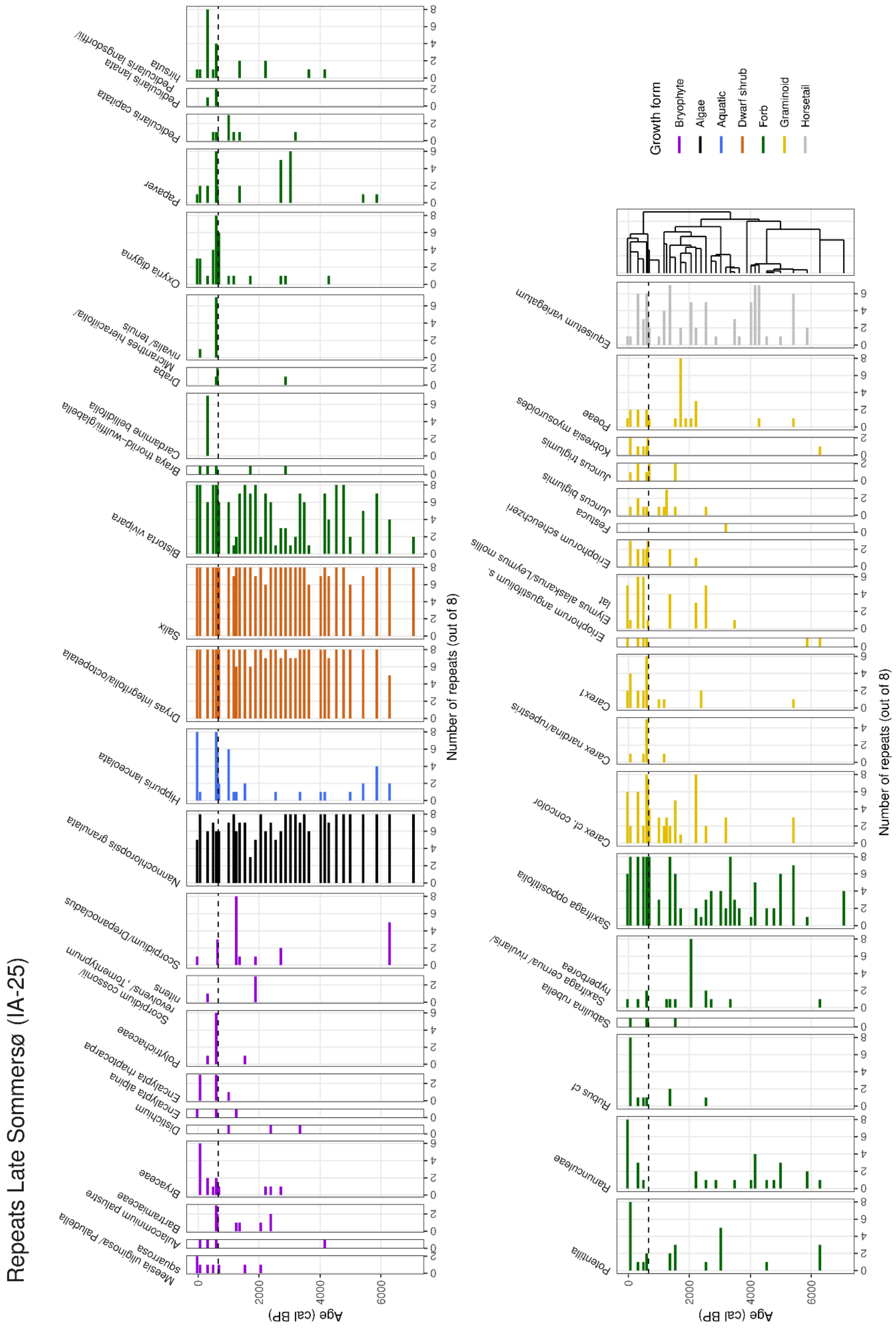

Repeats Late Sommersø (IA-25)
Age (cal BP)
Number of repeats (out of 8)
Growth form
Bryophyte
Algae
Aquatic
Dwarf shrub
Forb
Graminoid
Horsetail
Meesia uliginosa / Paludella squarrosa
Aulacomnium palustre
Bartramiaceae
Bryaceae
Distichium
Encalypta alpina
Encalypta rhaptocarpa
Polytrichaceae
Scorpidium cossonii/ revolvens / , Tomentypnum nitens
Scorpidium/Drepanocladus
Nannochloropsis granulata
Hippuris lanceolata
Dryas integrifolia/octopetala
Salix
Bistorta vivipara
Braya thorild−wulffii/glabella
Cardamine bellidifolia
Draba
Micranthes hieracifolia/ nivalis/ tenuis
Oxyria digyna
Papaver
Pedicularis capitata
Pedicularis lanata
Pedicularis hirsuta/ langsdorffii
Potentilla
Ranunculeae
Rubus cf
Sabulina rubella
Saxifraga cernua/ rivularis/ hyperborea
Saxifraga oppositifolia
Carex cf. concolor
Carex nardina/rupestris
Carex1
Eriophorum angustifolium s. lat
Elymus alaskanus/Leymus mollis
Eriophorum scheuchzeri
Festuca
Juncus biglumis
Juncus triglumis
Kobresia myosuroides
Poeae
Equisetum variegatum

b)

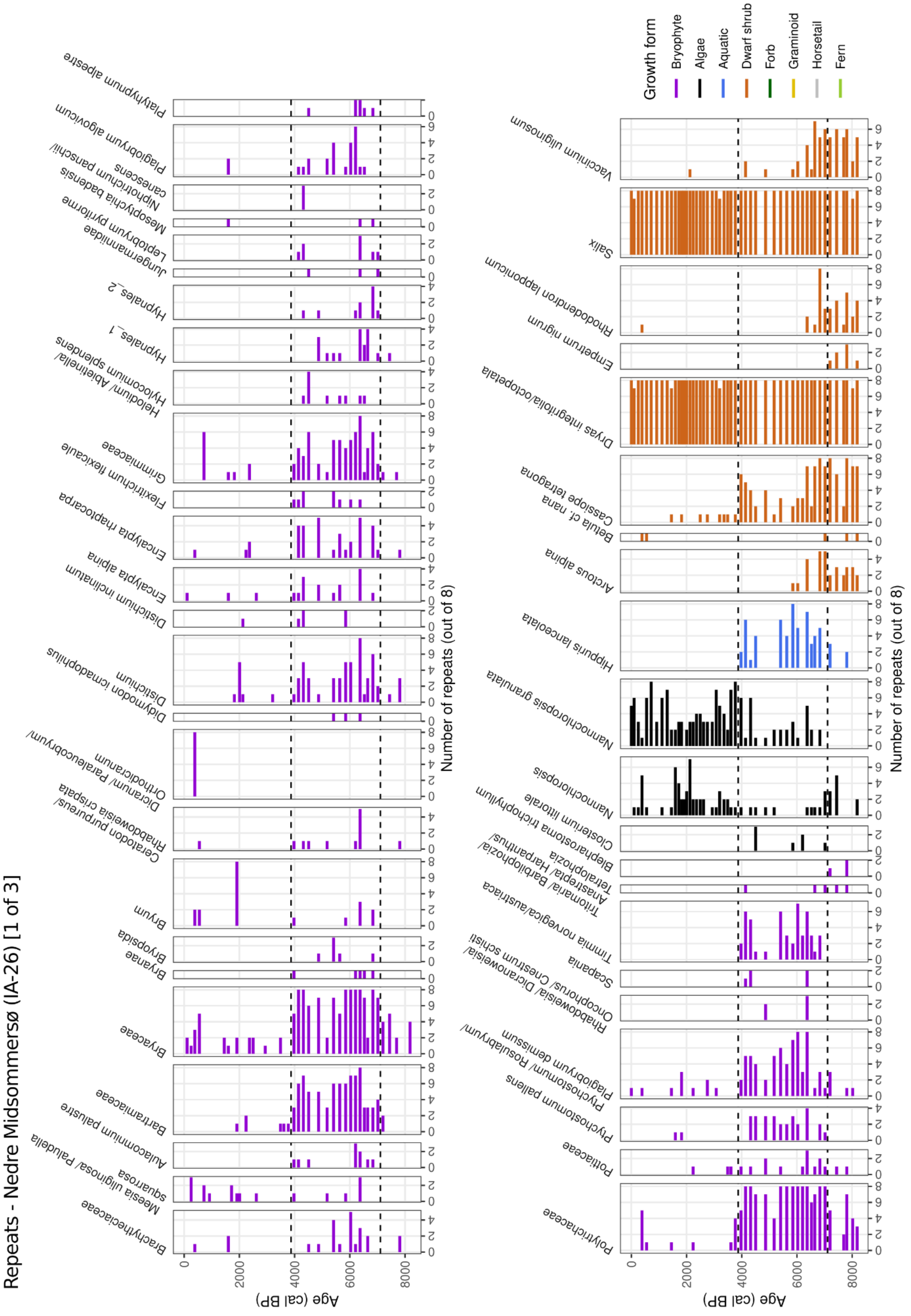
Repeats - Nedre Midsommersø (IA-26) [1 of 3]
Brachytheciaceae
Meesia uliginosa/ Paludella squarrosa
Aulacomnium palustre
Bartramiaceae
Bryaceae
Bryanae
Bryopsida
Bryum
Ceratodon purpureus/ Rhabdoweisia crispata
Dicranum/ Paraleucobryum/ Orthodicranum
Didymodon icmadophilus
Distichium
Distichium inclinatum
Encalypta alpina
Encalypta rhaptocarpa
Flexitrichum flexicaule
Grimmiaceae
Helodium/ Abietinella/ Hylocomium splendens
Hypnales_1
Hypnales_2
Jungermanniidae
Leptobryum pyriforme
Mesoptychia badensis
Niphotrichum panschii/ canescens
Plagiobryum algovicum
Platyhypnum alpestre
Polytrichaceae
Pottiaceae
Ptychostomum pallens
Ptychostomum/ Rosulabryum/ Plagiobryum demissum/
Rhabdoweisia/ Dicranoweisia/ Oncophorus/ Cnestrum schisti
Scapania
Timmia norvegica/austriaca
Tritomaria/ Barbilophozia/ Anastrepta/ Harpanthus/ Tetralophozia
Blepharostoma trichophyllum
Closterium littorale
Nannochloropsis
Nannochloropsis granulata
Hippuris lanceolata
Arctous alpina
Betula cf. nana
Cassiope tetragona
Dryas integrifolia/octopetala
Empetrum nigrum
Rhododendron lapponicum
Salix
Vaccinium uliginosum
Growth form
Bryophyte
Algae
Aquatic
Dwarf shrub
Forb
Graminoid
Horsetail
Fern
Age (cal BP)
Number of repeats (out of 8)

#### Repeats - Nedre Midsommersø (IA-26) [2 of 3]

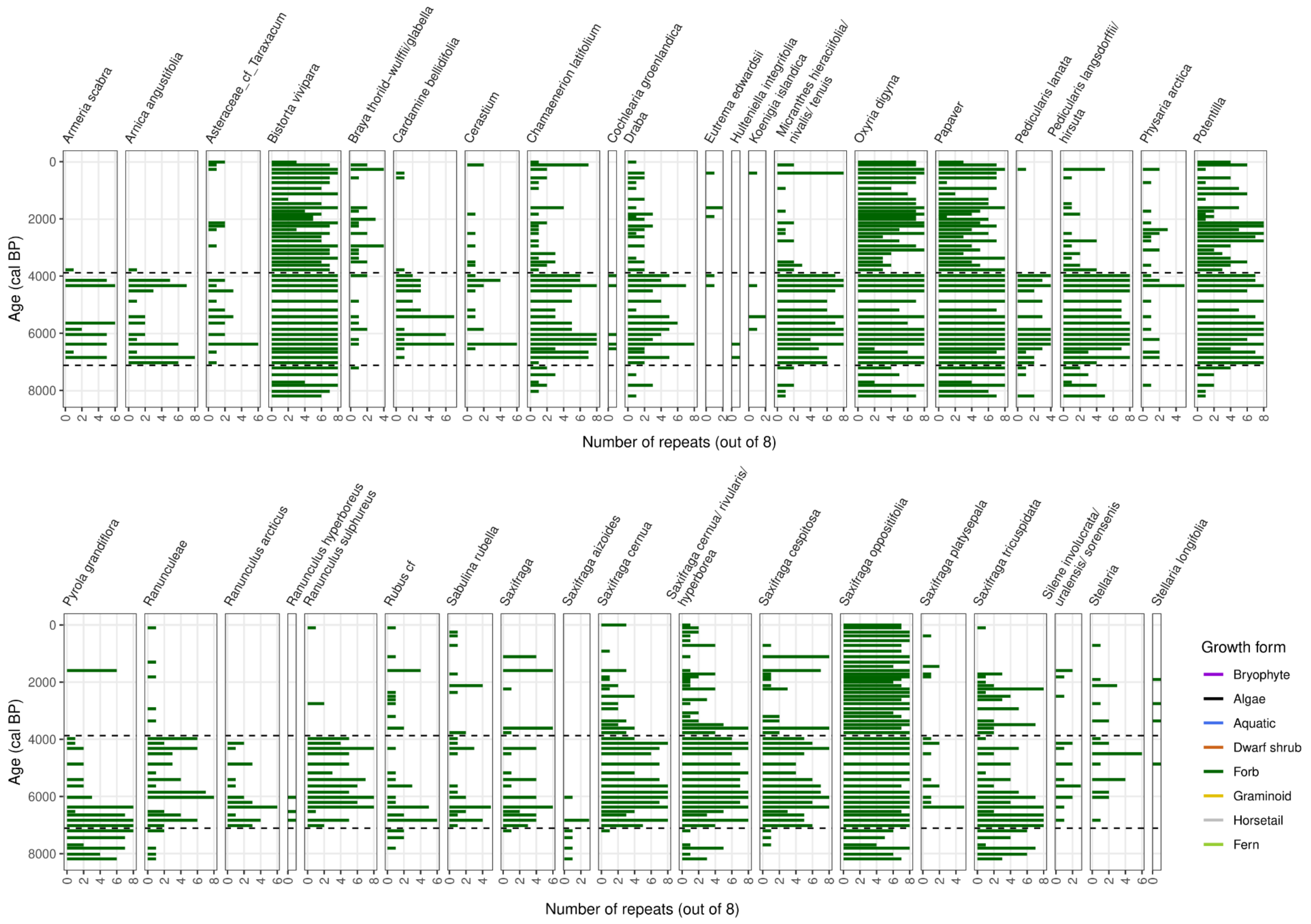

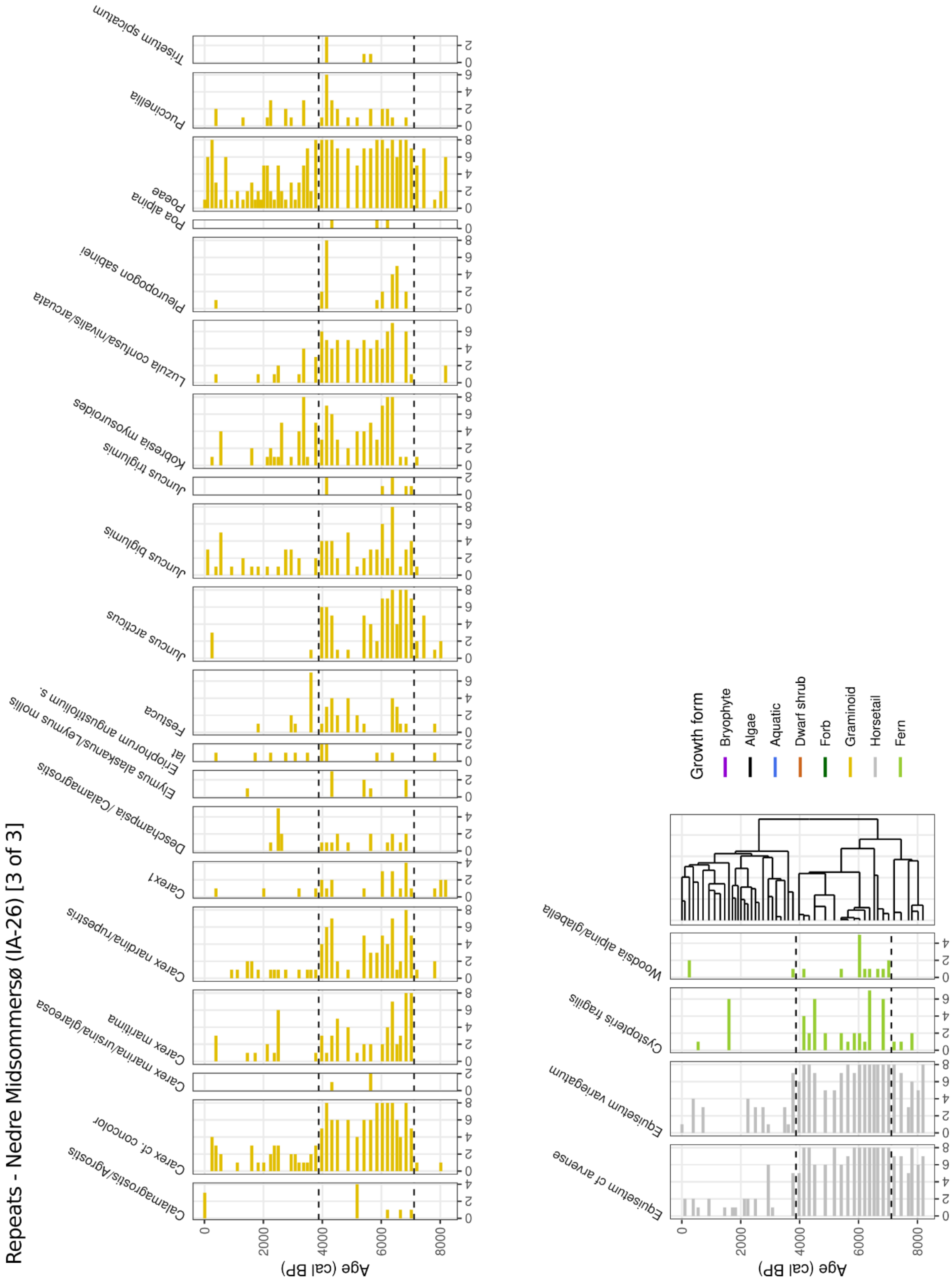
Repeats - Nedre Midsommersø (IA-26) [3 of 3]
Calamagrostis/Agrostis
Carex cf. concolor
Carex marina/ursina/glareosa
Carex maritima
Carex nardina/rupestris
Carex1
Deschampsia /Calamagrostis
Elymus alaskanus/Leymus mollis
Eriophorum angustifolium s. lat
Festuca
Juncus arcticus
Juncus biglumis
Juncus triglumis
Kobresia myosuroides
Luzula confusa/nivalis/arcuata
Pleuropogon sabinei
Poa alpina
Poeae
Puccinellia
Trisetum spicatum
Age (cal BP)
Equisetum cf arvense
Equisetum variegatum
Cystopteris fragilis
Woodsia alpina/glabella
Age (cal BP)
Growth form
Bryophyte
Algae
Aquatic
Dwarf shrub
Forb
Graminoid
Horsetail
Fern

c)

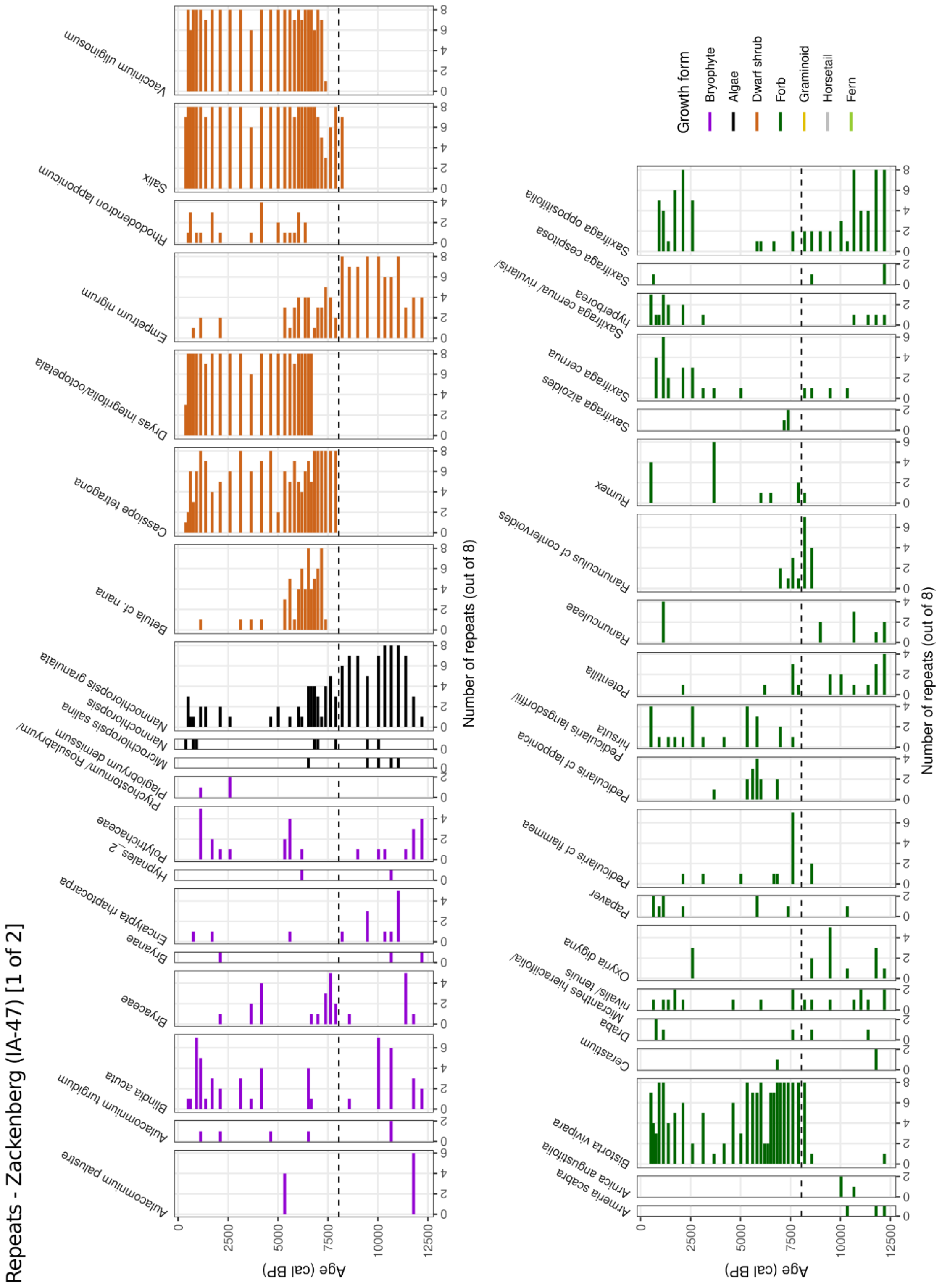

Repeats - Zackenberg (IA-47) [1 of 2]
Aulacomnium palustre
Aulacomnium turgidum
Blindia acuta
Bryaceae
Bryanae
Encalypta rhaptocarpa
Hypnales_2
Polytrichaceae
Plagiobryum demissum
Ptychostomum/ Rosulabryum/
Microchloropsis salina
Nannochloropsis
Nannochloropsis granulata
Betula cf. nana
Cassiope tetragona
Dryas integrifolia/octopetala
Empetrum nigrum
Rhododendron lapponicum
Salix
Vaccinium uliginosum
Armeria scabra
Arnica angustifolia
Bistorta vivipara
Cerastium
Draba
Micranthes hieraciifolia/
nivalis/ tenuis
Oxyria digyna
Papaver
Pedicularis cf flammea
Pedicularis cf lapponica
Pedicularis langsdorffii/
hirsuta
Potentilla
Ranunculaceae
Ranunculus cf confervoides
Rumex
Saxifraga aizoides
Saxifraga cernua
Saxifraga cernua/ rivularis/
hyperborea
Saxifraga cespitosa
Saxifraga oppositifolia
Age (cal BP)
Number of repeats (out of 8)
Growth form
Bryophyte
Algae
Dwarf shrub
Forb
Graminoid
Horsetail
Fern

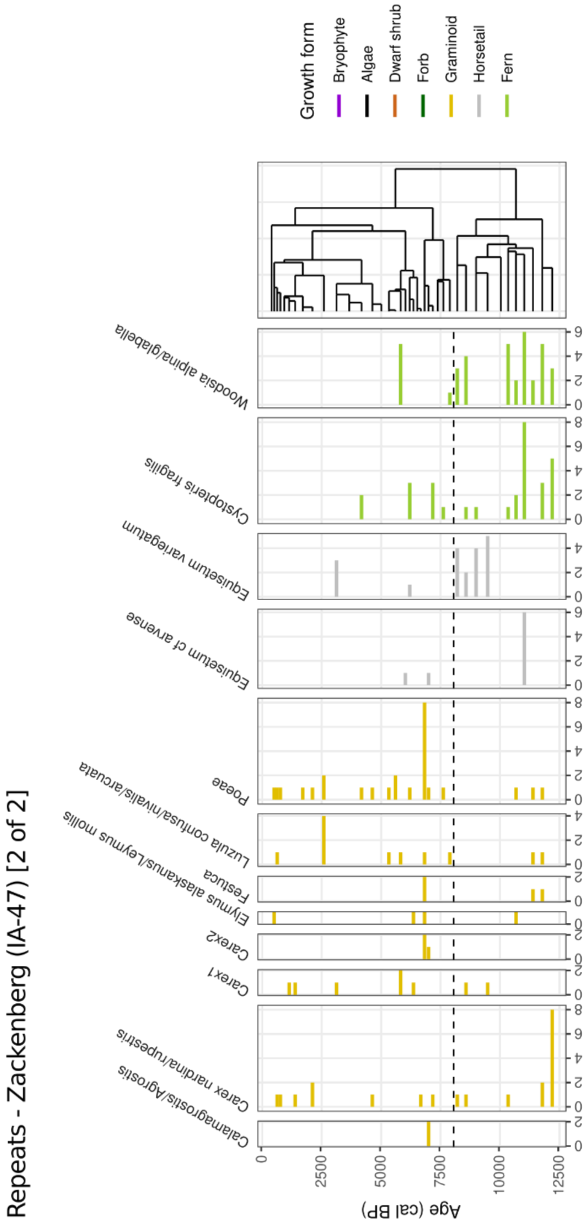


*Supplementary Fig. 3.1. The full metabarcoding plant record identified at a) Late Sommerso, b) Nedre Midtsommersø, and c) Zackenberg. The quantification is given as numbers of PCR repeats (maximum 8). CONISS zones are indicated by broken lines.*

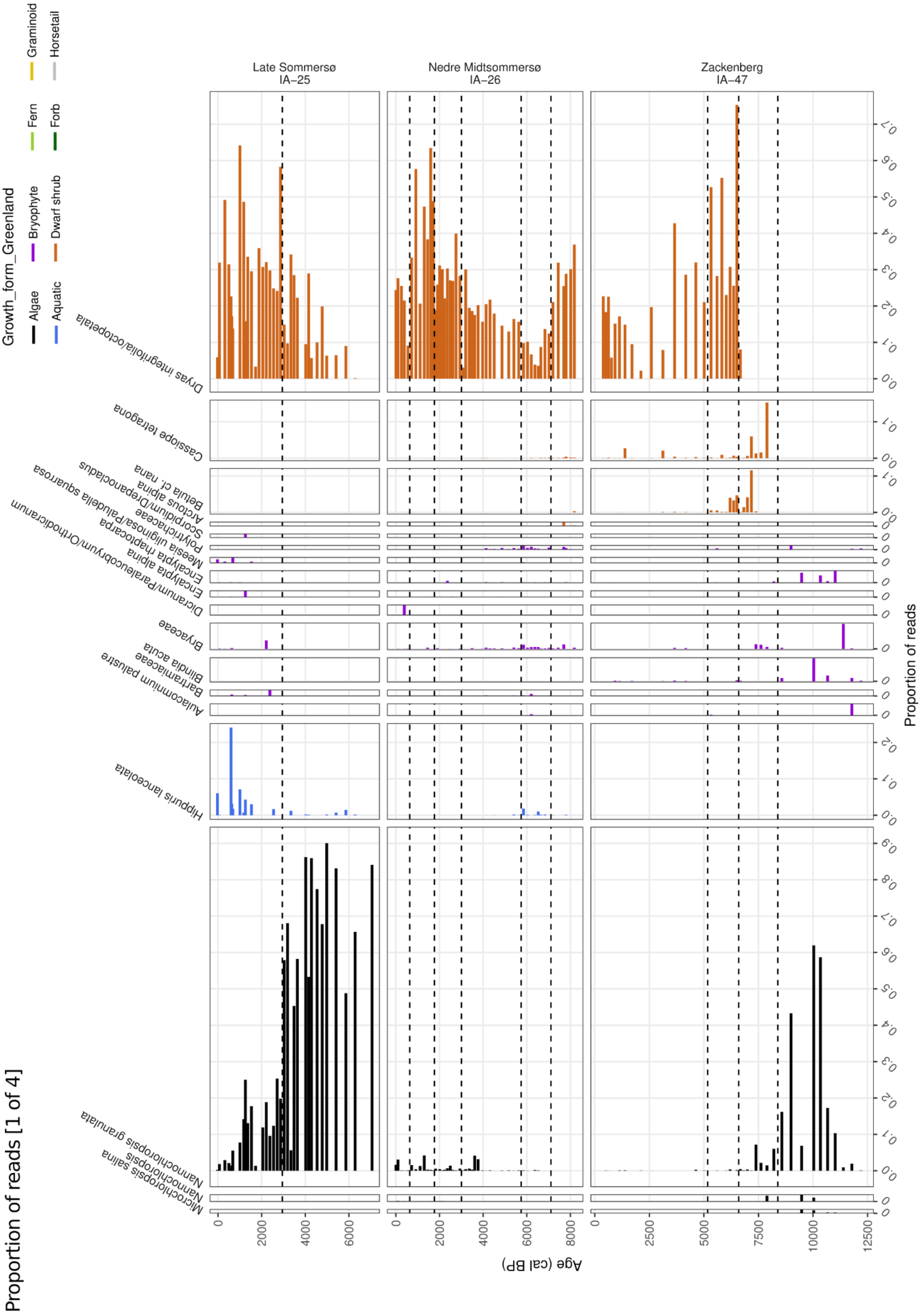
Proportion of reads [1 of 4]
Growth_form_Greenland
Algae
Aquatic
Bryophyte
Dwarf shrub
Fern
Forb
Graminoid
Horsetail
Late Sommersø
IA–25
Nedre Midtsommersø
IA–26
Zackenberg
IA–47
Dryas integrifolia/octopetala
Cassiope tetragona
Betula cf. nana
Bryaceae
Bartramiaceae
Hippuris lanceolata
Nannochloropsis granulata
Nannochloropsis salina
Microchloropsis salina
Age (cal BP)
Proportion of reads

Proportion of reads [2 of 4]

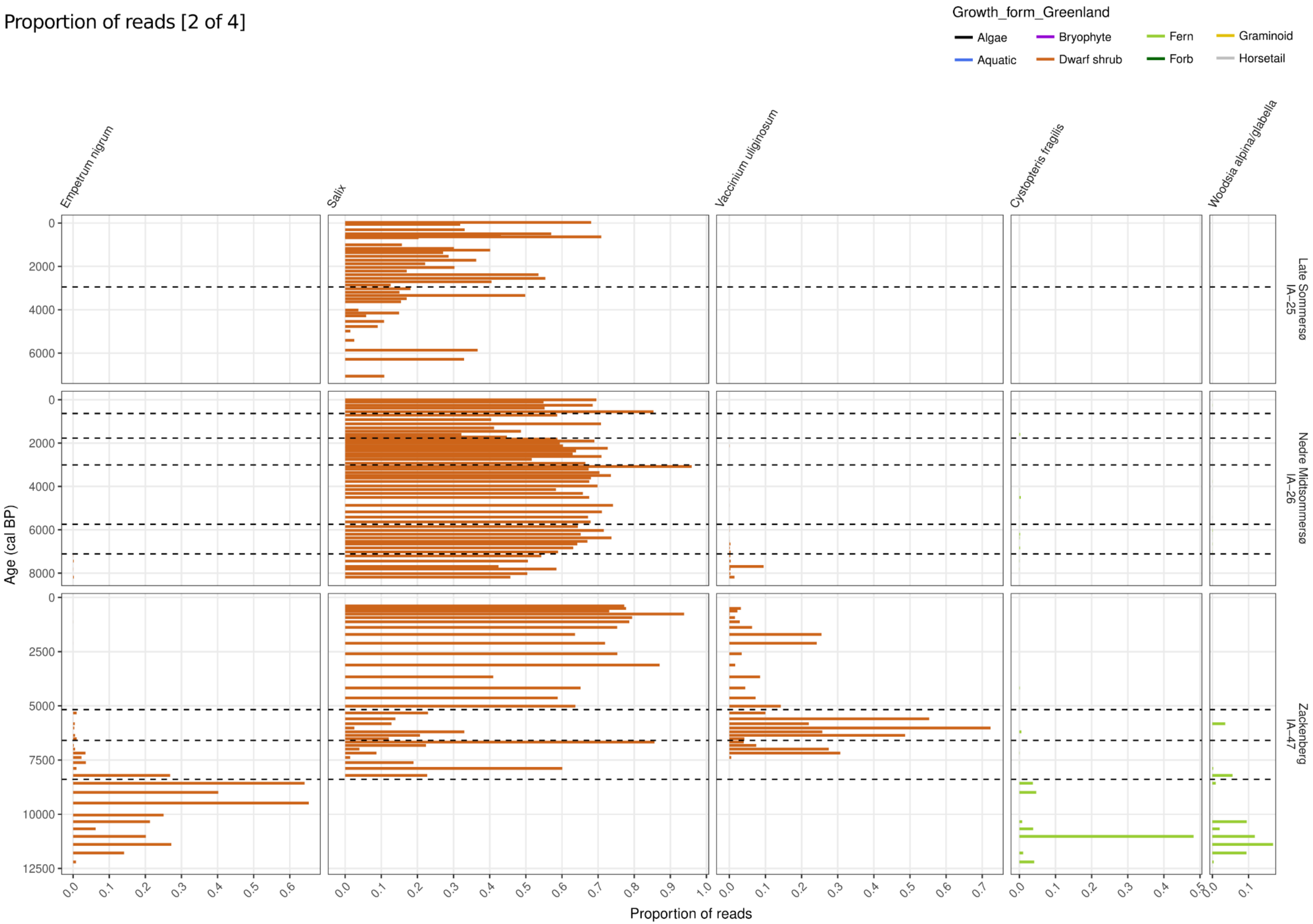

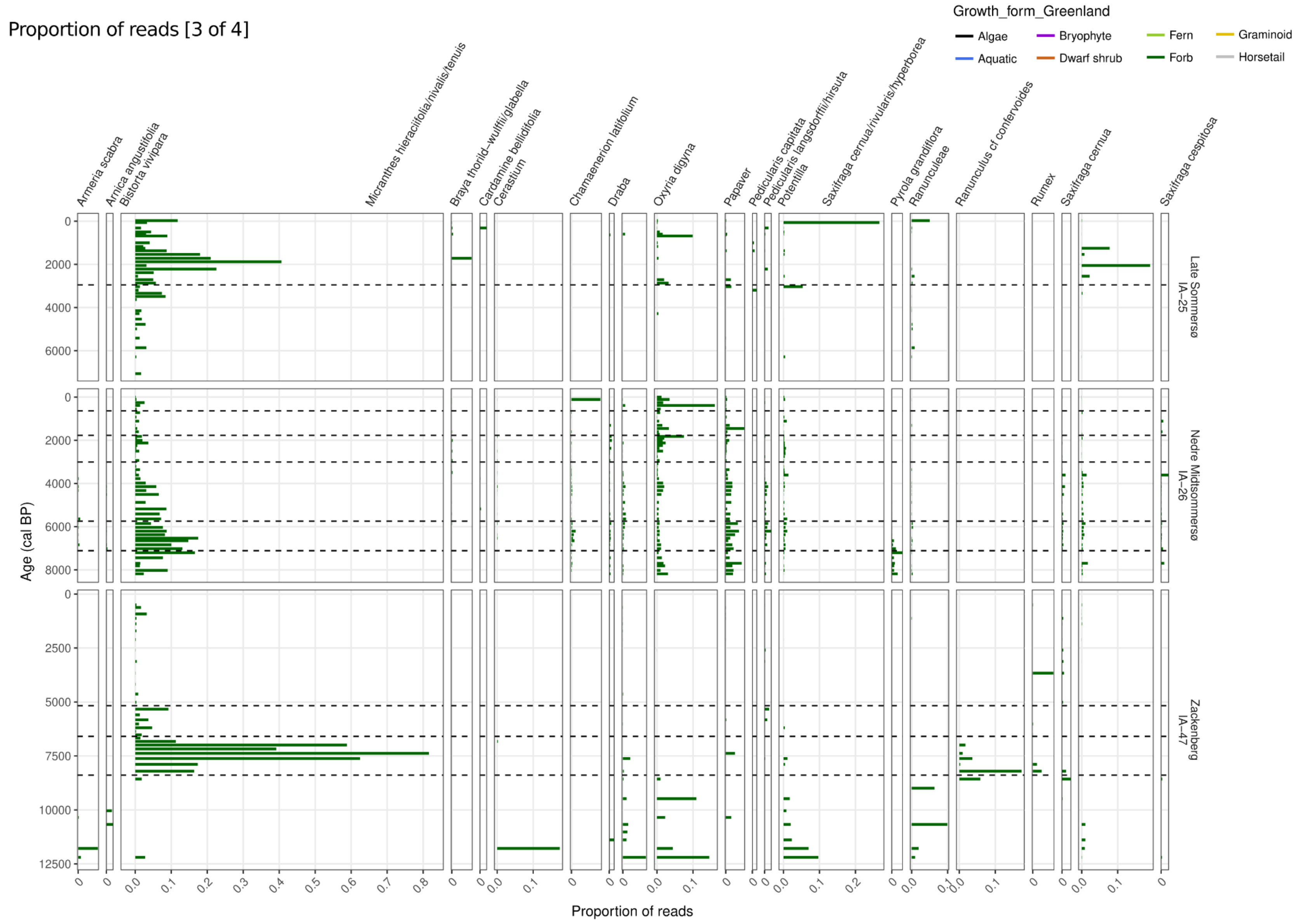

Proportion of reads [3 of 4]
Growth_form_Greenland
Algae
Aquatic
Bryophyte
Dwarf shrub
Fern
Forb
Graminoid
Horsetail
Armeria scabra
Arnica angustifolia
Bistorta vivipara
Micranthes hieraciifolia/nivalis/tenuis
Braya thorild–wulffii/glabella
Cardamine bellidifolia
Cerastium
Chamaenerion latifolium
Draba
Oxyria digyna
Papaver
Pedicularis capitata
Pedicularis langsdorffii/hirsuta
Potentilla
Saxifraga cernua/rivularis/hyperborea
Pyrola grandiflora
Ranunculeae
Ranunculus cf confervoides
Rumex
Saxifraga cernua
Saxifraga cespitosa
Late Sommersø
IA–25
Nedre Midtsommersø
IA–26
Zackenberg
IA–47
Age (cal BP)
Proportion of reads

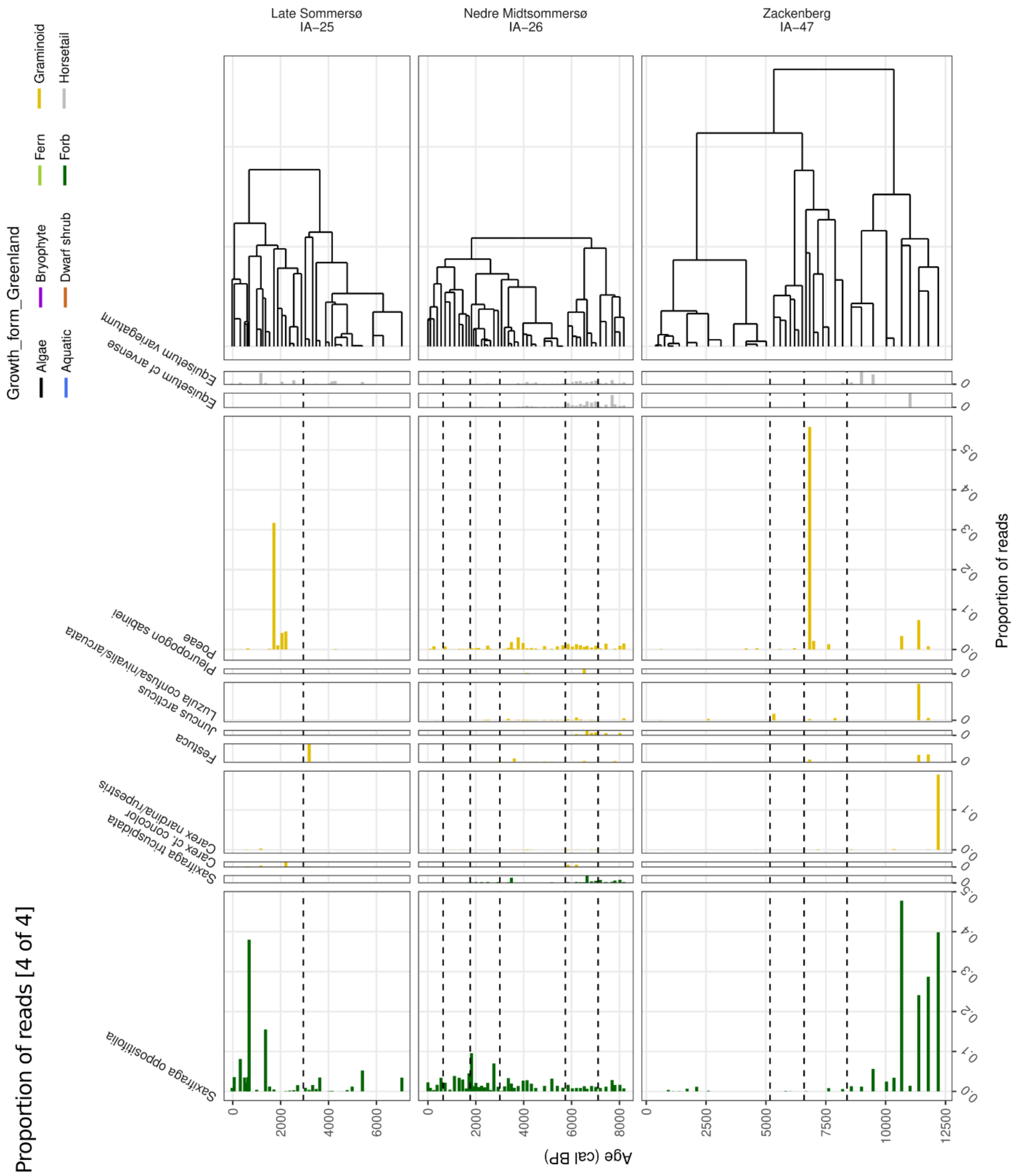


*Supplementary Fig. 3.2. Plant taxa recorded representing at least 1% of the total reads in one sample or more from Late Sommersø, Nedre Midtsommersø, and Zackenberg. The total number of reads is defined as all reads identified to native taxa (Supplementary Data 1). CONISS zones are indicated by broken lines.*

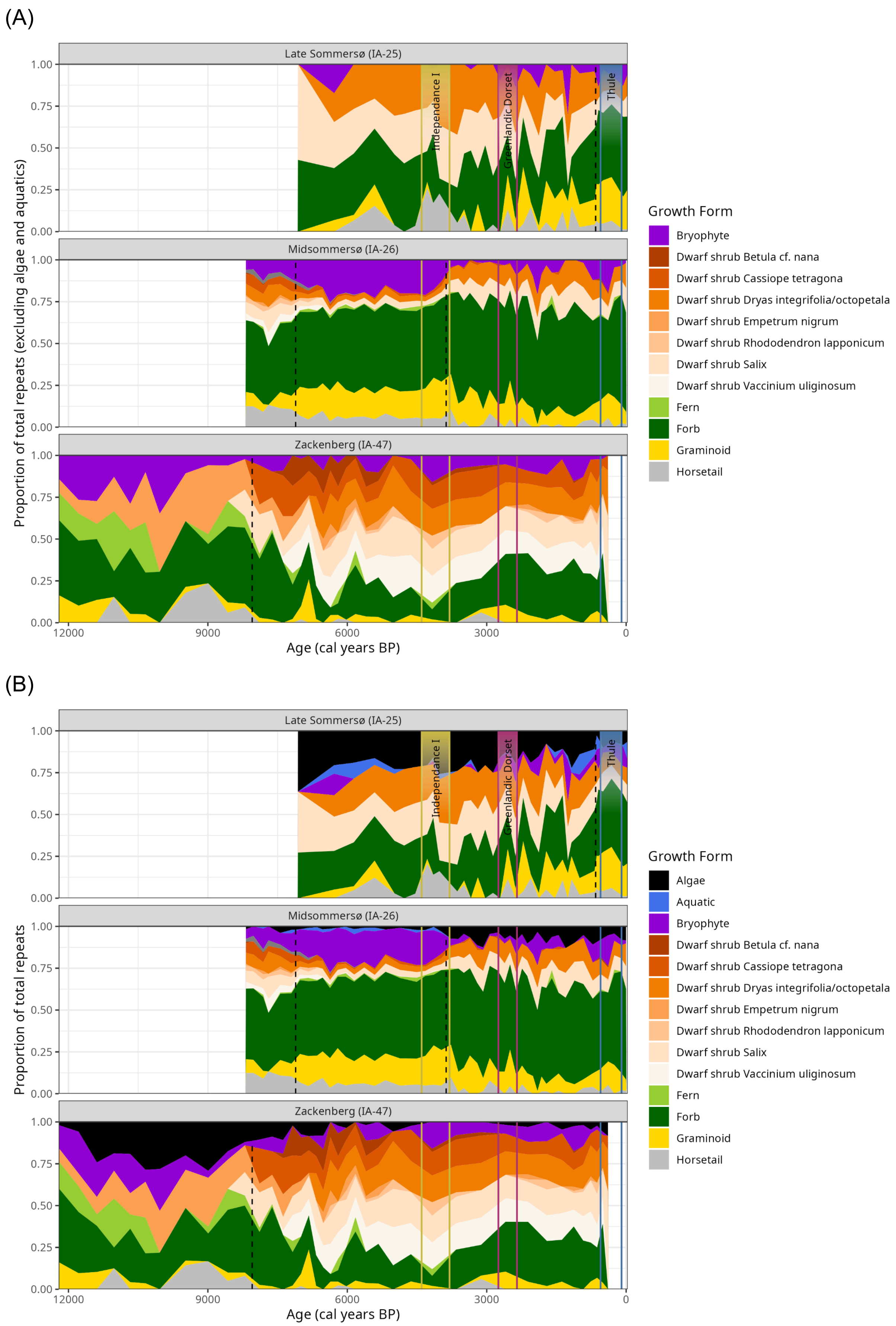

(A)
Late Sommersø (IA-25)
Midsommersø (IA-26)
Zackenberg (IA-47)
Proportion of total repeats (excluding algae and aquatics)
1.00
0.75
0.50
0.25
0.00
Independance I
Greenlandic Dorset
Thule
12000
9000
6000
3000
0
Age (cal years BP)
Growth Form
Bryophyte
Dwarf shrub Betula cf. nana
Dwarf shrub Cassiope tetragona
Dwarf shrub Dryas integrifolia/octopetala
Dwarf shrub Empetrum nigrum
Dwarf shrub Rhododendron lapponicum
Dwarf shrub Salix
Dwarf shrub Vaccinium uliginosum
Fern
Forb
Graminoid
Horsetail
(B)
Late Sommersø (IA-25)
Midsommersø (IA-26)
Zackenberg (IA-47)
Proportion of total repeats
Age (cal years BP)
Growth Form
Algae
Aquatic
Bryophyte
Dwarf shrub Betula cf. nana
Dwarf shrub Cassiope tetragona
Dwarf shrub Dryas integrifolia/octopetala
Dwarf shrub Empetrum nigrum
Dwarf shrub Rhododendron lapponicum
Dwarf shrub Salix
Dwarf shrub Vaccinium uliginosum
Fern
Forb
Graminoid
Horsetail

(C)

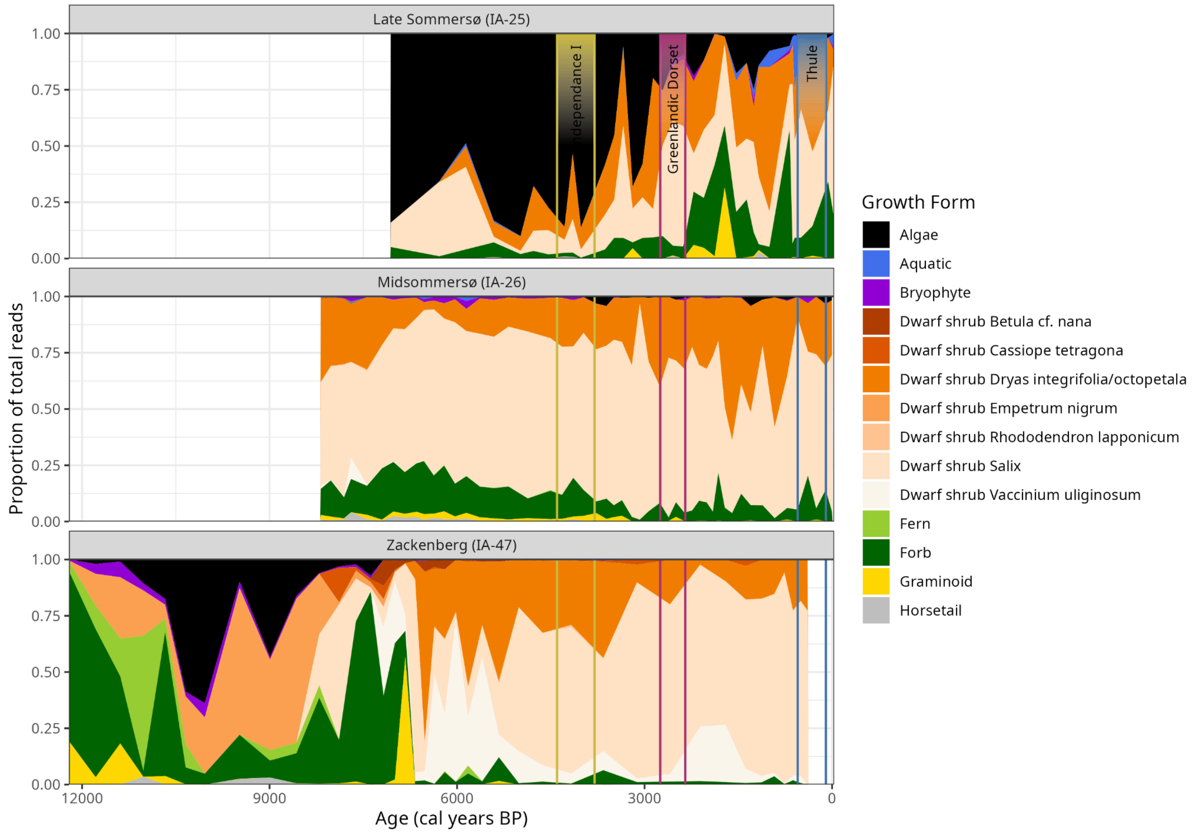


*Supplementary Fig. 3.3. Changes in plant composition over time. Change in proportion of PCR repeats identified to (A) terrestrial plants and (B) terrestrial and aquatic plants for Late Sommersø, Nedre Midtsommersø, and Zackenberg. (C) Change in proportion of total reads. CONISS zones are indicated by broken lines.*

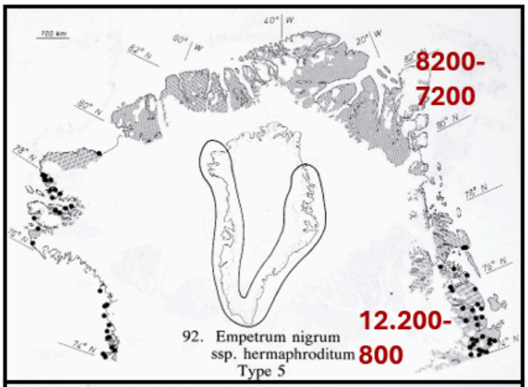


92. Empetrum nigrum ssp. hermaphroditum
Type 5

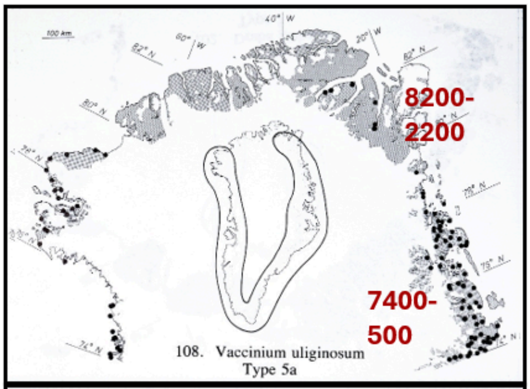


108. Vaccinium uliginosum
Type 5a

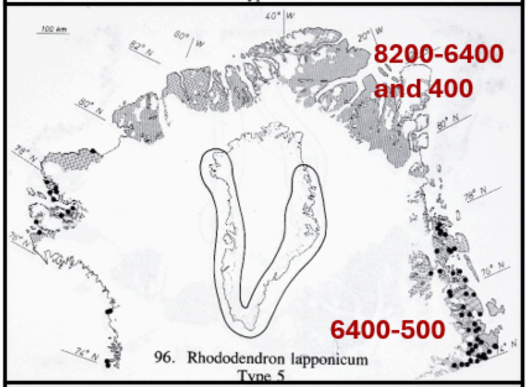


96. Rhododendron lapponicum
Type 5

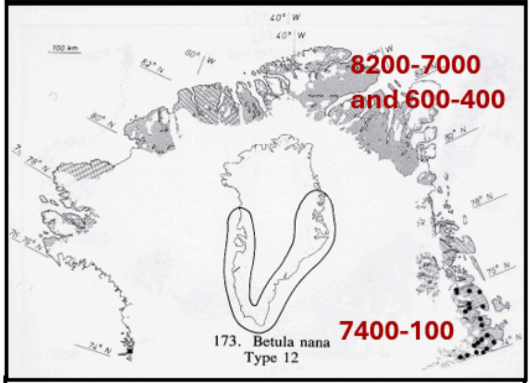


173. Betula nana
Type 12

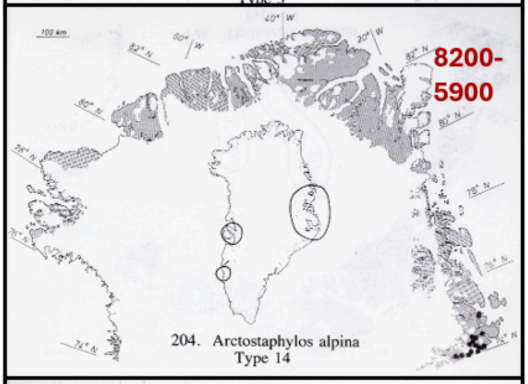


204. Arctostaphylos alpina
Type 14

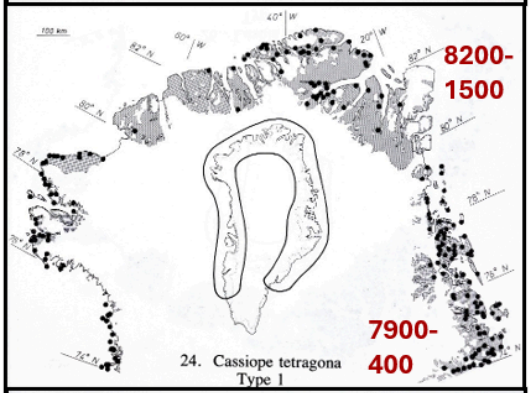


24. Cassiope tetragona
Type 1

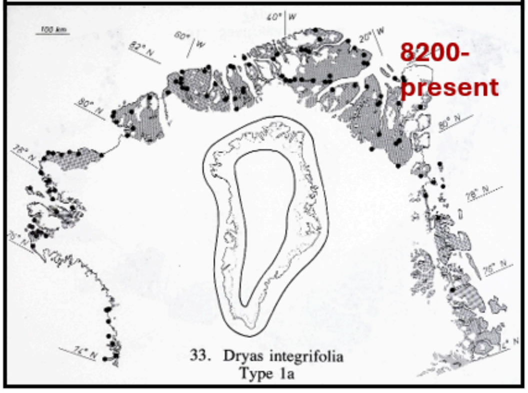


33. Dryas integrifolia
Type 1a

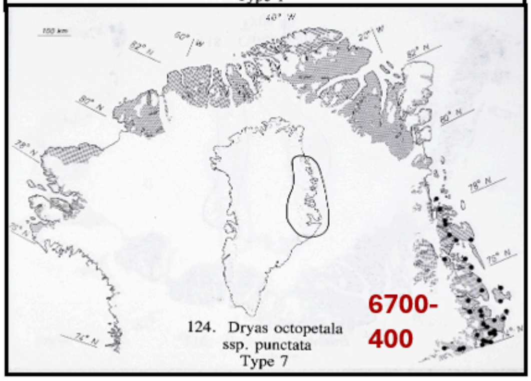


124. Dryas octopetala ssp. punctata
Type 7

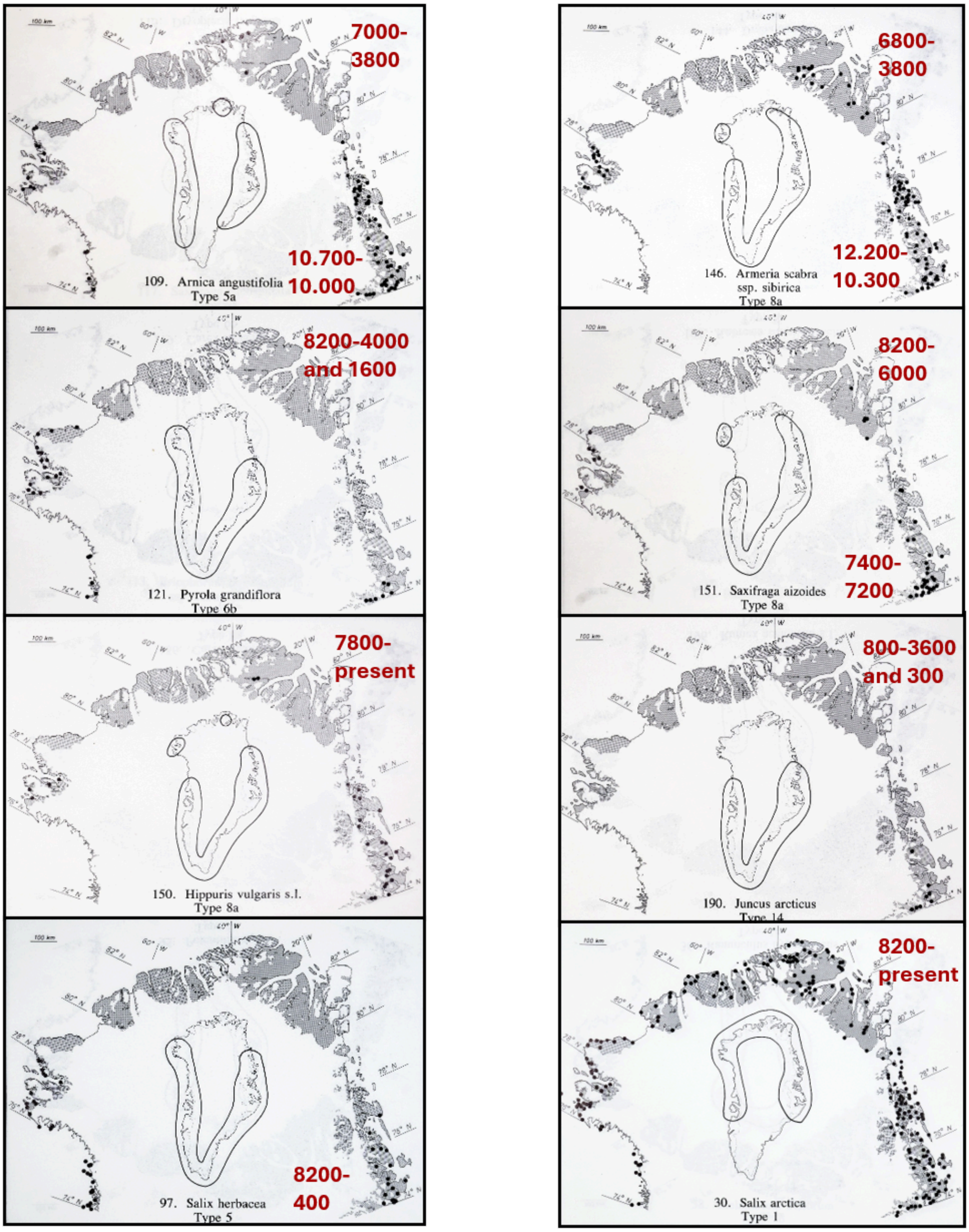


*Supplementary Fig. 3.4. Examples of current plant distribution and the first occurrences in or records from Wandel Dal and Zackenberg. The distribution type and maps are according to (5). Note that with the exception of* Salix, Dryas *and* Cassiope tetragona, *all examples are species with a clear northern thermal limit and only scattered distribution in the region today.*

### 4 The full animal metabarcoding record

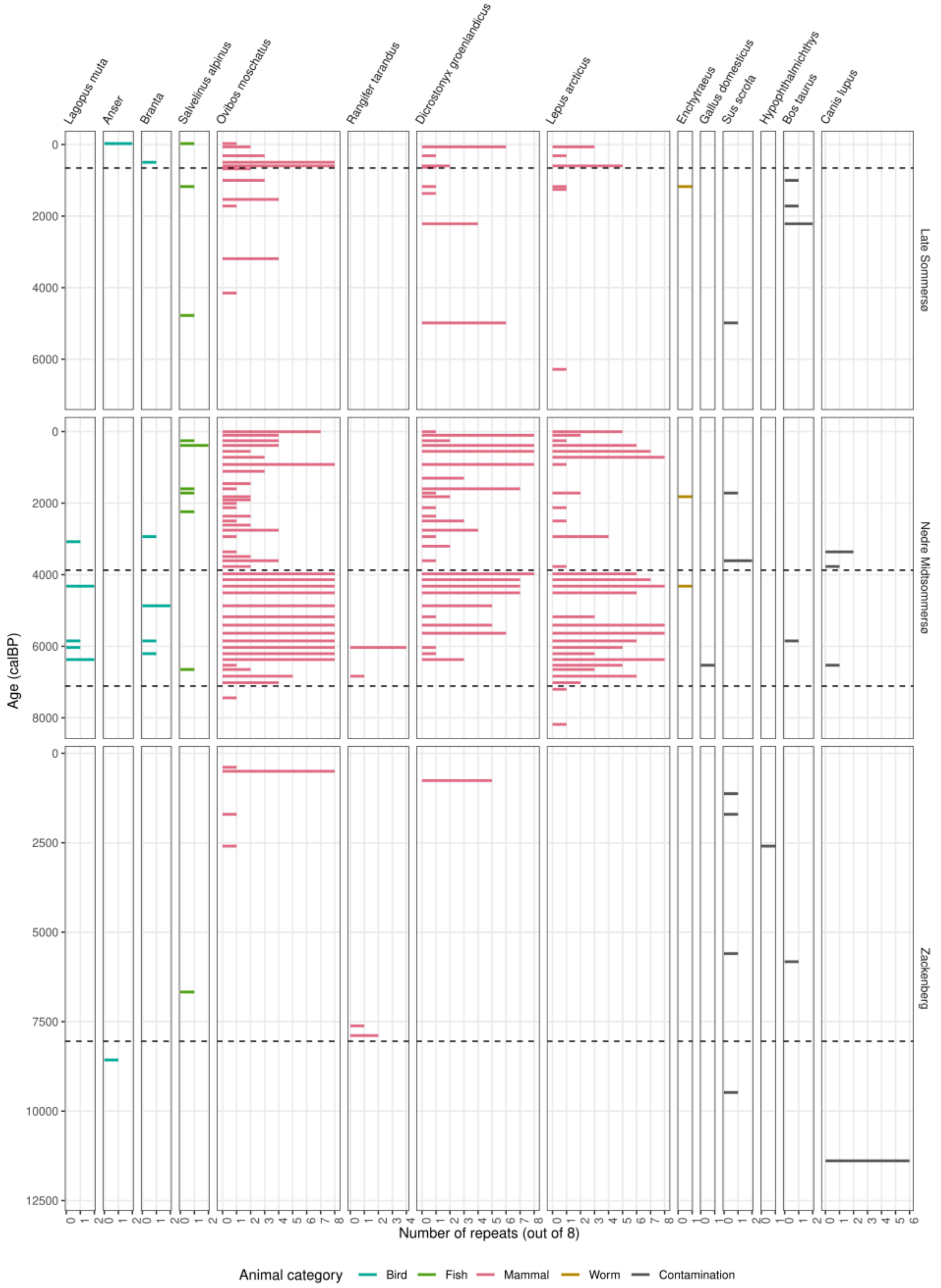


*Supplementary Fig. 4.1. The full animal record identified at Late Sommersø (top), Nedre Midtsommersø (middle), and Zackenberg (bottom). The quantification is given as numbers of PCR repeats (maximum 8). The figures include contaminants (black bars) including* Canis lupus *likely representing dogs.*

### 5 The shotgun sequencing results

The shotgun analyses of 32 samples and 6 negative controls resulted in a total of 15.4 billion raw reads, with 41,216,213 and 111,257 reads matching plants and animals of putative ancient origin, respectively (Supplementary Data 2).

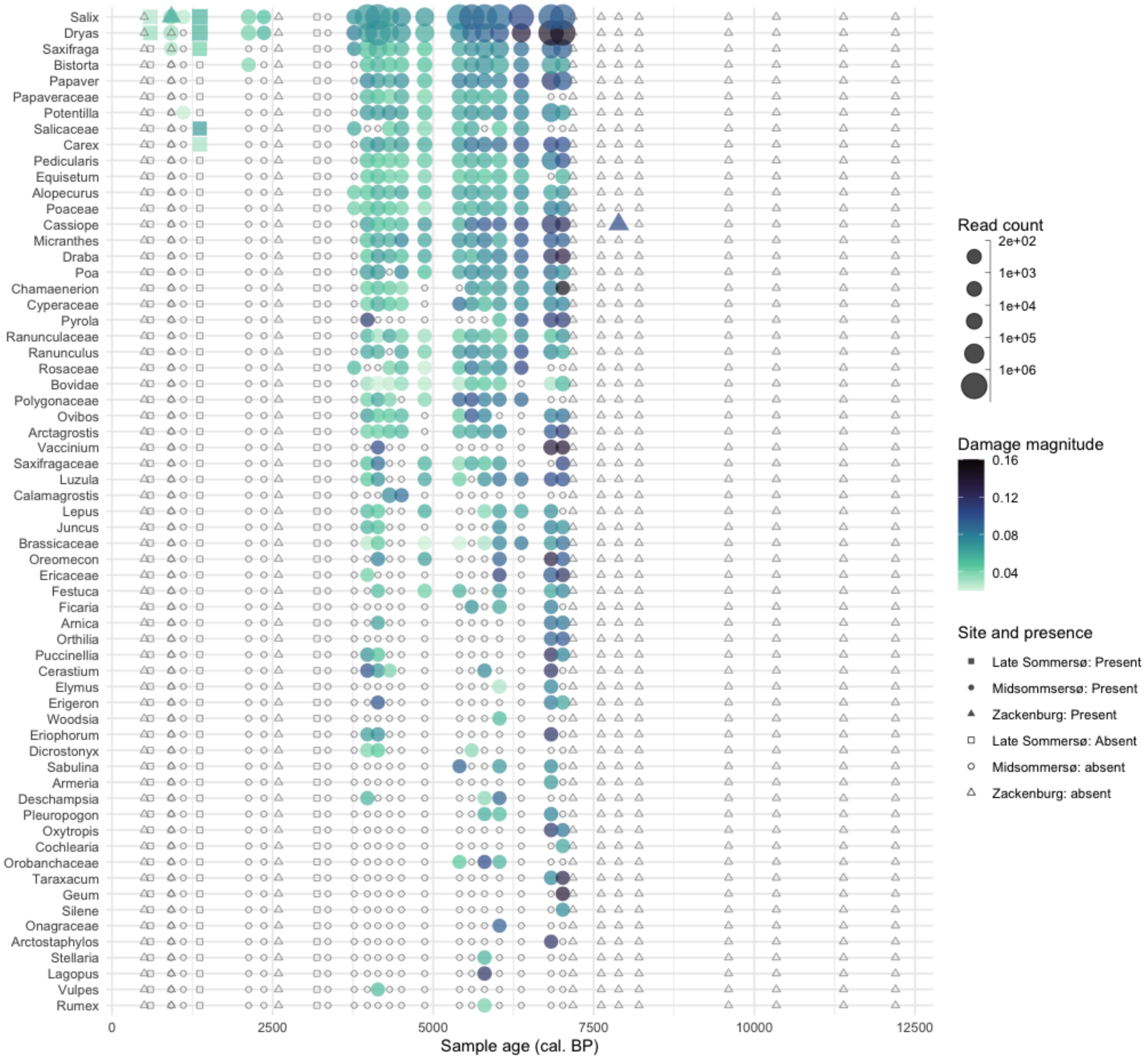


*Fig. 5.1 Taxa identified in shotgun sequencing determined to be ancient in origin. Solid shapes indicate that the taxon was present while hollow shapes indicate the taxon was not detected at that site with the most stringent filtering including significant damage pattern. The full record of taxa identified in the shotgun data are presented in Supplementary Data 2.*

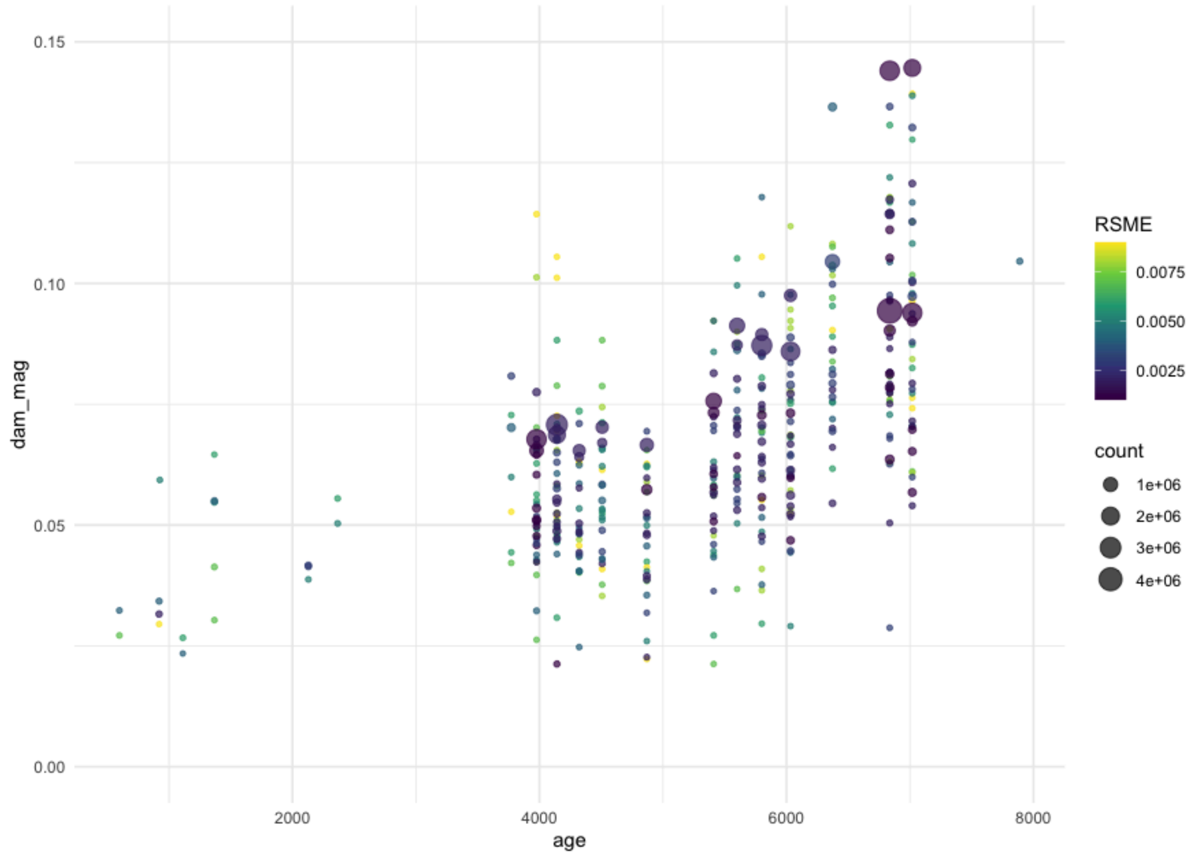


*Fig. 5.2 Damage pattern magnitude of taxa in relation to sample age. Each dot represents the damage pattern observed of a single taxon in each sample across the three sites. Root square mean error (RSME) from pyDamage's GLM is represented by the color of each point.*

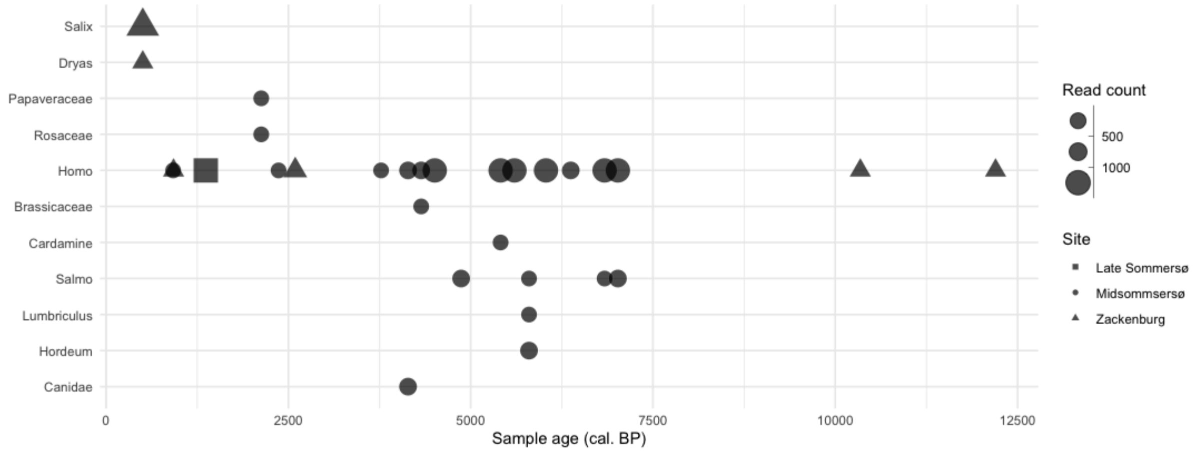


*Fig. 5.3. Taxa identified in shotgun sequencing without a significant damage pattern. Note that the sample that has Canidae (IA-26-2-28) is not the same as had Canis in metabarcoding: IA-47_DNA38, IA-26-1_SED23, IA-26-1_SED24, and IA-26-3_SED37. Only the latter was analysed for shotgun, but no Canin passed the filters applied.*

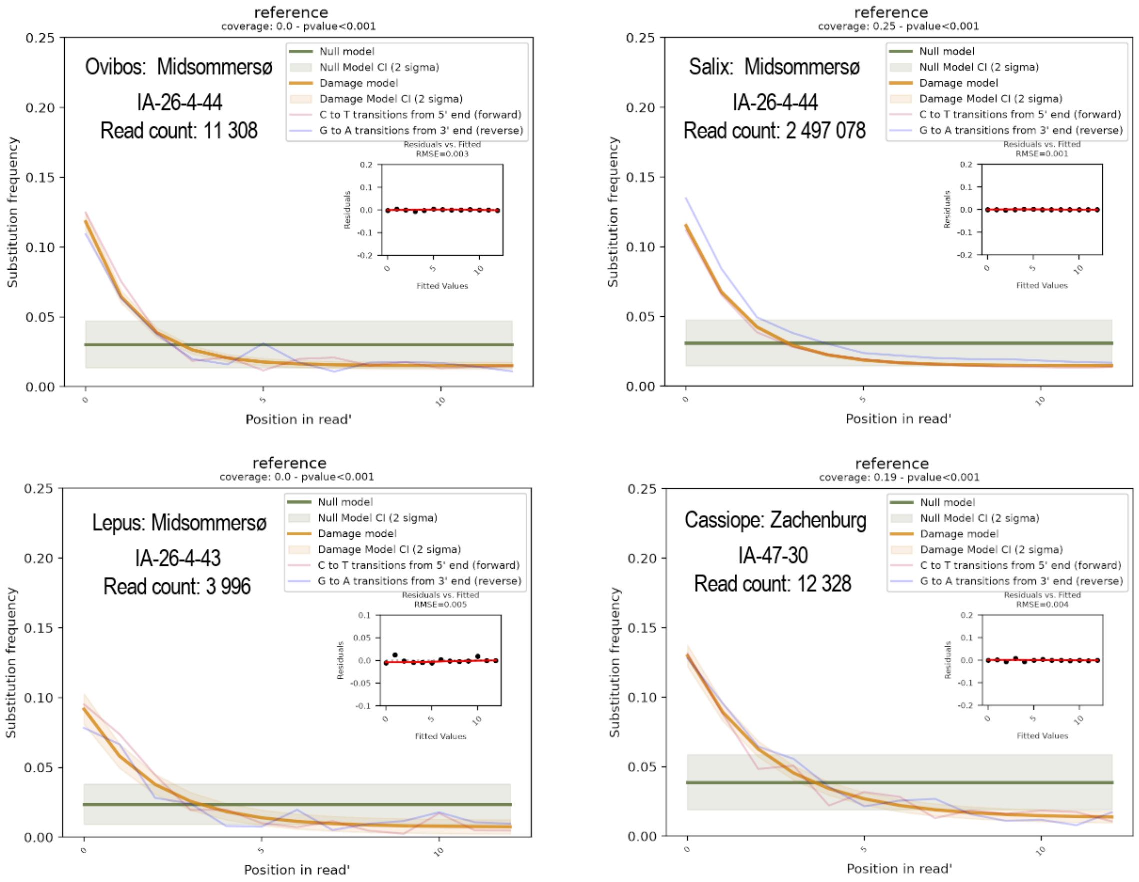


*Fig. 5.4. Damage plots generated by pyDamage for the first occurrence of four taxa;* Ovibos, Salix, Lepus*, and* Cassiope*.*

### 6. Comparison of metabarcoding and shotgun metagenomics data

As the number of samples analysed for metabarcoding (n=138) is much higher than the number of samples analysed for shotgun (n=23), we compare the methods per taxa rather than per sample. This also takes into account that detection of taxa that deposit low quantity of DNA in the environment, like rare and scattered plants (6, 7) and most mammals (8) are not likely to be detected in every sample.

#### Vascular plants

For some taxa, the identification was on different taxonomic level so we cannot say for sure if the for example the different species of grasses identified in the metabarcoding correspond to the grasses identified in shotgun data. Many taxa were potentially identified at higher taxonomic resolution in metabarcoding than shotgun, for example species of *Saxifraga* and *Carex* (Supplementary Table S7). *Betula* cf. *nana*, *Hippuris lanceaolata, Hulteniella integrifolia* and *Cystopteris fragilis* were detected but not passing our filtering control of the shotgun data. Two taxa, *Erigeron* and *Geum* were identified in the shotgun data (Supplementary Data 2) but not in metabarcoding. While the former is common in the region today, the latter only has a single occurrence in NE Greenland and that is south-east of Wandel Dal and otherwise only south Greenland (5, 9). Two further taxa, *Ficara* and

*Oxytropis* were also identified in the shotgun data but are not likely for the region (but the family Ranuculaceae is so *Ficaria* might represent a mis-identification).

##### Animals

All mammals that were frequently observed in the metabarcoding data like muskox, northern collard lemming and arctic hare, were also observed in shotgun data (Supplementary Table 8, Supplementary Data 2). One mammal, *Vulpes*, presumably representing the Arctic fox, was found only in one sample of shotgun data. On the other hand, reindeer were found in two samples of metabarcoding, confirmed by dated macrofossils from the region but missing from the shotgun data probably because of scattered local populations combined with few samples from the relevant period. For other vertebrates, one of the tentative three bird taxa (rock ptarmigan) was confirmed in shotgun analyses and the only fish identified was only in metabarcoding. Further, none invertebrate passed the filtering criteria in shotgun probably due to lack of reference genomes.

#### References


1. N. L. Balascio, *et al.*, Midsommersø records the Holocene glacial history of Wandel Dal, Inutoqqat Nunaat (Peary Land) northern Greenland. *Quat. Sci. Rev.* **378**, 109874 (2026).

2. M. Blaauw, J. A. Christen, Flexible paleoclimate age-depth models using an autoregressive gamma process. *Bayesian Anal.* **6**, 457–474 (2011).

3. M. Blaauw, Methods and code for “classical” age-modelling of radiocarbon sequences. *Quat. Geochronol.* **5**, 512–518 (2010).

4. D. P. Rijal, *et al.*, Sedimentary ancient DNA shows terrestrial plant richness continuously increased over the Holocene in northern Fennoscandia. *Science Advances* **7**, eabf9557 (2021).

5. C. Bay, A phytogeographical study of vascular plants of northern Greenland - north of 74° northern latitude. *Meddelelser om Grønland, Bioscience* **36**, 1–102 (1992).

6. I. G. Alsos, *et al.*, Plant DNA metabarcoding of lake sediments: How does it represent the contemporary vegetation. *PLoS One* **13**, e0195403 (2018).

7. T. G. Ataman, Y. Lammers, I. G. Alsos, D. P. Rijal, A. G. Brown, Sedimentary DNA from lake depocenters maximizes detection of catchment vegetation. *Commun. Earth Environ.* **6**, 1–10 (2025).

8. T. J. Murchie, C. Giguet-Covex, P. D. Heintzman, V. Slon, Y. Wang, “Terrestrial fauna and hominin DNA from sedimentary archives” in *Developments in Paleoenvironmental Research*, (Springer International Publishing, 2023), pp. 299–378.

9. F. Rune, *Wild flowers of Greenland* (Gyldenlund Naturforlag, 2011).
